# The Drosophila OSC Genome: A Resource for Studies of Transposon and piRNA Biology

**DOI:** 10.1101/2025.10.09.681131

**Authors:** Dominik Handler, Julius Brennecke

## Abstract

Accurate genome assemblies are critical for understanding small RNA-mediated genome defense. In animals, the PIWI-interacting RNA (piRNA) pathway protects genome integrity by silencing transposable elements. Studying how piRNAs are generated and how they guide heterochromatin formation requires complete reconstruction of genomic piRNA source loci and detailed transposon maps.

Here, we present a high-quality *de novo* genome assembly of *Drosophila melanogaster* ovarian somatic cells (OSCs), a widely used cell line that recapitulates nuclear piRNA biology. The OSC genome differs substantially from the reference genome, with major differences in transposon content and piRNA cluster composition. Our assembly resolves the 700 kb *flamenco* locus, the primary piRNA cluster in OSCs, and provides a genome-wide transposon map.

Using this resource, we characterize piRNA source loci, reveal how piRNA cluster composition determines transposon-derived piRNA profiles, and clarify the widespread impact of the nuclear piRNA pathway on heterochromatin. Finally, we provide an open platform for integrating user-generated datasets with the OSC genome, creating a community resource for studying transposon control and piRNA biology.

## INTRODUCTION

Transposable elements are mobile genetic elements that occupy large portions of eukaryotic genomes and threaten genome stability [1–3]. In plants, fungi, and animals, their activity is restrained by small RNA-based silencing systems, which use Argonaute proteins to repress transposon expression at transcriptional or post-transcriptional levels [4–6]. The specificity of these pathways depends on the small RNAs bound by Argonaute proteins, which are produced predominantly from transposon-rich regions of the genome.

Because many small RNAs involved in genome defense originate from repetitive or strain-specific transposon loci, they often map to multiple genomic locations. This complicates the reconstruction of their genomic origins and obscures their relationships with active transposons. Accurate and complete genome assemblies therefore greatly facilitate the study of small RNA origins, diversity, and targets.

In animals, genome defense is mediated primarily by the PIWI-interacting RNA (piRNA) pathway, a conserved mechanism that operates from sponges to mammals. It acts mainly in the gonads, where it silences transposons in germ cells and in surrounding somatic support cells [7–9].

The fruit fly *Drosophila melanogaster* is a leading model for studying the piRNA pathway, supported by its rich genetic toolkit and long-standing role in transposon research [10, 11]. A key experimental system is the ovarian somatic cell (OSC) line, one of the very few animal cell lines that maintains a functional, transposon-repressing piRNA pathway [12, 13].

The piRNA pathway in OSCs mirrors that of somatic follicle cells in the *Drosophila* ovary and depends on a single nuclear PIWI-clade Argonaute protein, Piwi [13–16]. Unlike germ cells, OSCs lack the cytoplasmic PIWI proteins Aubergine and Ago3 and therefore do not employ the ping-pong amplification cycle of sense and antisense piRNAs [17, 18]. Instead, single-stranded piRNA precursors enriched in transposon antisense sequences are processed on the mitochondrial surface into phased 22-32 nt piRNAs that are loaded into Piwi [7–9].

Once bound to piRNAs, Piwi is imported into the nucleus [19], where it recognizes nascent transposon transcripts through sequence complementarity. Target engagement triggers assembly of the Piwi* complex—comprising Piwi, Asterix/Gtsf1, and Maelstrom—which recruits heterochromatin factors and enforces transcriptional silencing [20–30].

The simplicity and completeness of the OSC system (one Argonaute, one piRNA biogenesis pathway, one mode of silencing) make OSCs an ideal model for dissecting primary piRNA biogenesis and piRNA-guided heterochromatin formation. However, despite their extensive use, the genomic framework of OSCs remains incompletely characterized.

Studies of piRNA biology in flies and OSCs have thus far relied on the *Drosophila melanogaster* reference genome [17, 31, 32]. The most recent assembly, dm6, which also resolves large portions of repeat-rich heterochromatin, has been crucial for identifying and characterizing genomic piRNA source loci (piRNA clusters). However, the reference genome represents a fly strain that differs markedly from most laboratory strains, including those used to establish OSCs [33]. In addition, cell line immortalization often involves extensive genomic rearrangements and bursts of transposon activity, generating new insertions that are absent in the reference genome. Consistent with this, short-read sequencing of OSC DNA revealed a distinct transposon landscape that diverges markedly from that of dm6 [24, 33]. These differences complicate the accurate mapping of piRNAs, the annotation of piRNA clusters, and the interpretation of chromatin profiling data, thereby limiting the precision of OSC-based studies of piRNA biology.

Here, we present a high-quality *de novo* assembly of the OSC genome. The assembly achieves scaffold continuity comparable to the *Drosophila melanogaster* reference genome and resolves all major piRNA source loci, including a complete reconstruction of the rapidly evolving *flamenco* cluster. In addition, it provides a comprehensive, genome-wide map of transposon insertions and reports on structural variants and single nucleotide polymorphisms, allowing the genome-wide design of effective siRNAs and guide RNAs. Together, these features establish a precise genomic framework for studying transposon control in OSCs and for interpreting piRNA and chromatin datasets at nucleotide resolution. To ensure broad accessibility, we provide the OSC assembly in standard formats and as an interactive UCSC Genome Browser hub, enabling seamless integration with user-generated data.

## RESULTS

### High-quality *de novo* assembly of the OSC genome

The *Drosophila melanogaster* ovarian somatic cell (OSC) line was established in 2009 as a stable derivative of the original mixed fGS/OSS culture, which itself originated from dissected *bam* mutant ovaries [12, 13, 33]. We obtained the OSC line from the Siomi laboratory in 2010 and used an early passage to isolate high-molecular-weight genomic DNA for sequencing. Oxford Nanopore long reads, and Illumina short reads provided ∼100x and ∼50x coverage, respectively, ensuring sufficient depth for a highly contiguous assembly.

To assemble the OSC genome, we developed a custom multi-step pipeline optimized to recover transposable element sequences (Figure 1A). Incorporation of Hi-C chromosome conformation capture data further improved scaffold continuity and chromosome-level organization. A particular challenge was the non-isogenic nature of the OSC genome, which reflects the mixed genetic background of the founder fly stock. Instead of generating separate haplotype genomes, we generated a consensus genome in which heterozygous transposon insertions and structural variants were retained by default and heterozygous SNPs were randomly chosen but fully reported.

**Figure 1.**
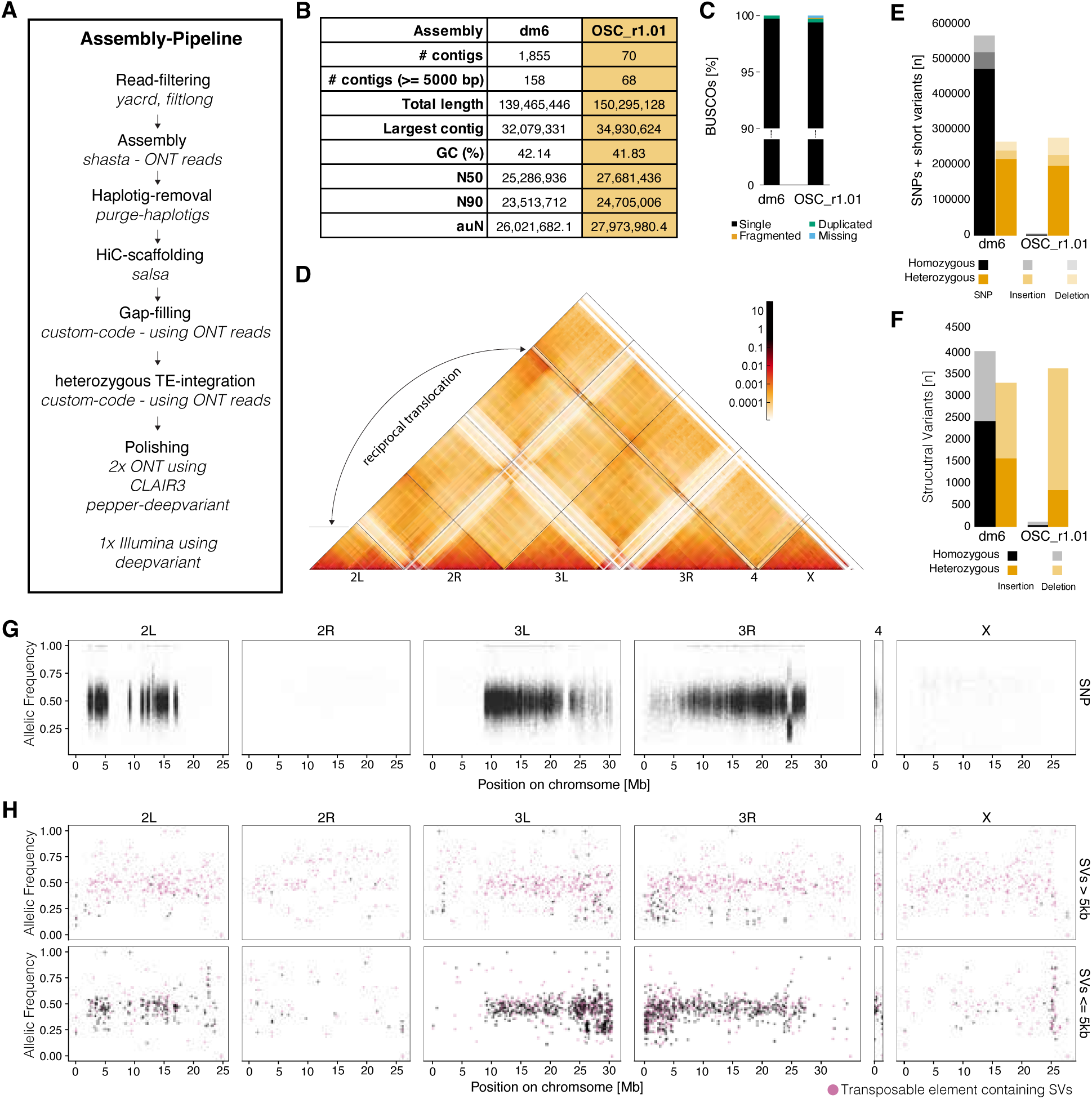
A highly accurate reference genome for OSC cells. **(A)** Assembly workflow integrating Oxford Nanopore long reads, Illumina short reads, and Hi-C data. **(B)** Assembly statistics of the dm6 reference genome and the OSC_r1.01 genome. **(C)** BUSCO completeness comparison between dm6 and OSC_r1.01 assemblies. **(D)** Hi-C contact map of the OSC_r1.01 genome showing a heterozygous translocation of distal 2L to 3R. **(E)** Homozygous and heterozygous SNPs and small variants identified by DeepVariant from Illumina reads obtained from OSC genomic DNA aligned to the dm6 or the OSC_r1.01 genome. **(F)** Homozygous and heterozygous structural variants (SVs) detected by Sniffles from Nanopore reads obtained from OSC genomic DNA aligned to the dm6 or the OSC_r1.01 genome. **(G)** Genome-wide SNP allele frequencies across chromosomes. **(H)** Genome-wide distribution of short (<5 kb, top) and long (=>5 kb, bottom) SV frequencies across chromosomes.

The final assembly (OSC_r1.01) spans ∼150 Mb and surpasses the current *Drosophila melanogaster* reference genome (dm6) across most quality metrics (Figure 1B, figure-supplement 1A). Even when scaffolds are broken into contigs, continuity remains higher than in dm6 (figure-supplement 1B). Benchmarking Universal Single-Copy Orthologs (BUSCO) analysis confirmed that the OSC genome is also highly complete, comparable in quality to the reference genome (Figure 1C). Although highly contiguous, the assembly does not yet extend telomere-to-telomere. As in other *Drosophila* assemblies, extremely repetitive regions—including telomeric transposon arrays, the histone gene cluster, rDNA repeats, and centromeres—remain only partially resolved.

To assess the structural integrity of our assembly, we mapped Hi-C reads obtained from OSCs onto the assembled OSC genome (Figure 1D). This analysis confirmed the overall assembly quality and revealed a reciprocal translocation of the distal ∼10 Mb of chromosome 2L with the end of chromosome 3R (figure-supplement 1C). Long nanopore reads confirmed this translocation on one homolog and located the breakpoint between *CG33298* and *Oatp30B* on chromosome 2L (figure-supplement 1D).

At the nucleotide level, we benchmarked the OSC genome by calling single nucleotide polymorphisms (SNPs) as well as short variants smaller than 50 bp (Figure 1E) and structural variants above 50 bp (Figure 1F). As expected, re-mapping OSC-derived Illumina and Nanopore reads to the OSC assembly yielded almost no homozygous aberrations. In comparison, mapping the OSC reads to the dm6 genome resulted in >500,000 homozygous differences, highlighting considerable differences between the two genomes. On average, the OSC genome differs from dm6 by ∼ 6 SNPs per kb, with important implications for experimental design, particularly when selecting effective siRNAs or CRISPR guide RNAs. For instance, the gene encoding the piRNA biogenesis factor SoYb harbors 85 homozygous nucleotide differences across its 4431 coding bases.

We next examined allelic variation to assess heterozygosity of the OSC cell line. In agreement with the mixed genetic background of the founder fly strain, heterozygous variants were abundant. We detected >250,000 heterozygous SNPs, reflecting extensive heterozygosity of the OSC genome. Notably, these variants were not evenly distributed: plotting SNP density along the chromosome arms revealed extensive ‘SNP deserts’, where the average SNP density dropped from ∼5 per kb to a background of <0.1 (Figure 1G). The SNP deserts correspond to previously described loss-of-heterozygosity (LOH) regions, thought to have arisen from segmental deletions or copy-neutral mitotic recombination events during cell line immortalization [33].

Genome-wide mapping of short and long structural variants showed that short variants were also largely absent from the LOH regions, whereas long variants were not (Figure 1H). Consistent with previous reports [33], many long variants correspond to transposable-element insertions, indicating that new transposition events occurred after the LOH episodes that shaped the immortalized OSC genome.

In summary, we report a high-quality assembly of the OSC genome with extensive continuity, high completeness, and detailed insights into heterozygosity and structural variation. This resource establishes a robust foundation for studies of transposon biology and piRNA-mediated genome defense.

### Annotation and genome browser integration of the OSC genome

To make the OSC genome broadly usable, we comprehensively annotated both genes and repeats (Figure 2A). Gene models from FlyBase [34] were aligned to the OSC genome, and transposable elements were annotated using RepeatMasker supplemented with a curated library of *Drosophila melanogaster* transposon consensus sequences [35]. Together, these annotations provide a detailed map of the gene and transposon landscape of the OSC genome.

**Figure 2.**
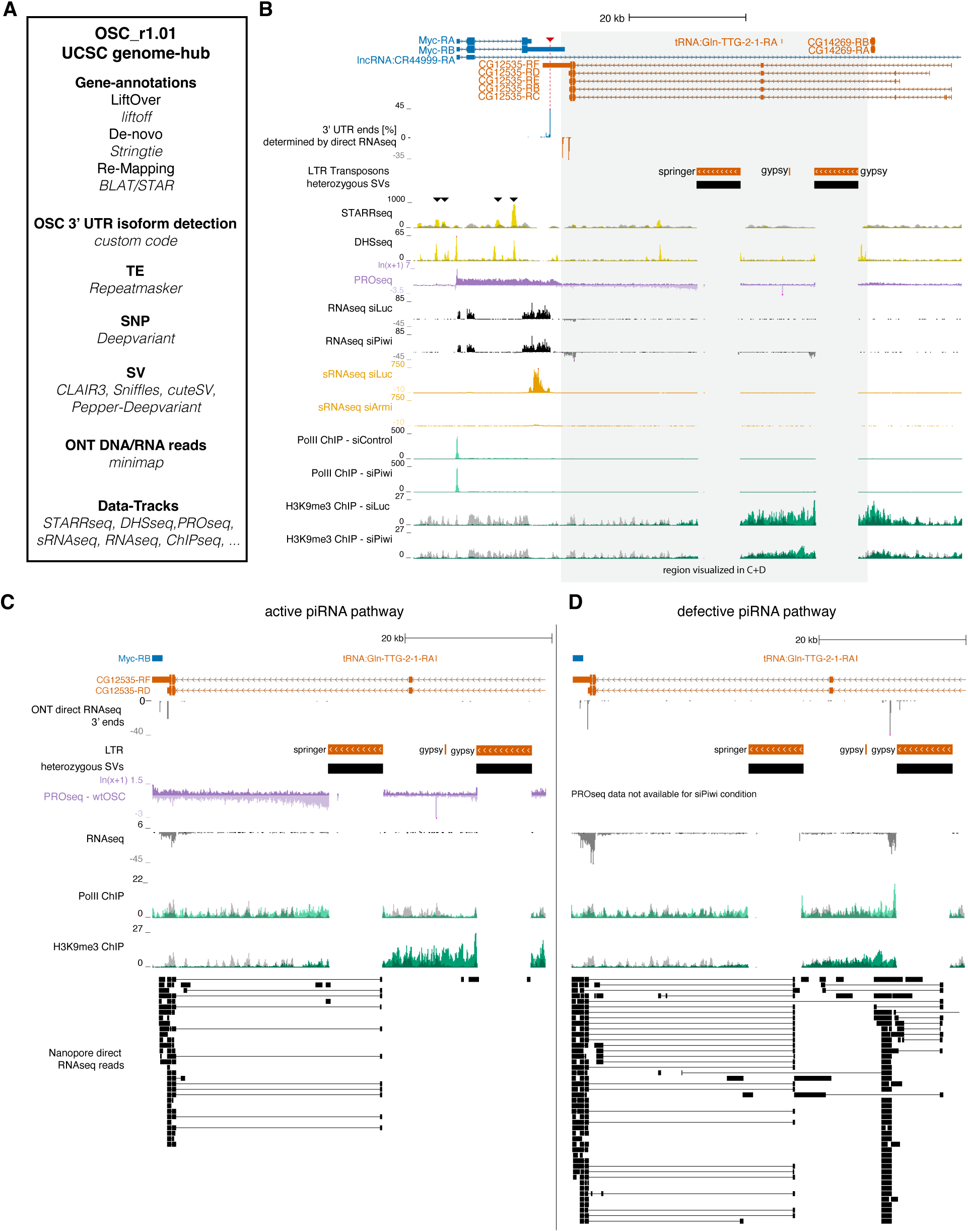
An accessible genome browser provides insights into the piRNA-transposon interplay. **(A)** Overview of datasets and genome annotations available in the OSC UCSC genome browser session with tools used for track generation or data type indicated. **(B)** Genome browser view of the *Myc* and *CG12535* loci. The extent of a novel 3′ UTR isoform of the *Myc* transcript identified by Nanopore direct RNA-seq 3′ end quantification is marked by a red arow. Individual detected genic 3′ ends are shown as a percentage of all detected 3′ ends for each gene. (STARR, DHS, ChIP, PRO and RNA signal are shown as coverage per million reads, small RNA coverage normalized to 1M miRNA reads, data is displayed as the average of 3 replicates. PRO-seq signal is displayed as ln(x + 1)-transformed coverage). ChIP-seq is shown as an overlay of ChIP (green) and ChIP input (grey). STARR and DHS-seq are shown as an overlay of experiment (yellow) and input (grey). STARR-seq peaks in them *Myc* locus are indicated with black arrows. (**C-D**) Genome browser views of the *CG12535* locus containing heterozygous *springer* and *gypsy* LTR retrotransposon insertions. Shown in addition to (B) are ONT direct RNA-seq 3′ ends from knockdowns (normalized to 1 million aligned reads). (**C**) Wild-type cells with an active piRNA pathway. (**D**) Piwi-depleted cells with a defective piRNA pathway (representative subset of Nanopore direct RNA-seq reads shown).

To enrich and validate these annotations, we integrated a wide range of published and newly generated sequencing datasets (supplementary table 1). We assembled a broad panel of datasets covering transcriptome activity (poly(A)-selected and rRNA-depleted RNA-seq, small RNA-seq, direct long read RNA-seq), transcriptional dynamics (PRO-seq, Pol II ChIP-seq), chromatin state (ChIP-seq for H3K9me3, DNase-seq for chromatin accessibility) and enhancer activity (STARR-seq for developmental and house-keeping genes). To maximize accessibility, we organized all annotations and functional datasets into a dedicated UCSC genome browser hub (figure-supplement 2A) [36]. This hub provides an interactive platform where users can explore the OSC genome at high resolution, compare functional data sets with FlyBase annotations, and integrate their own datasets.

To illustrate the utility of the OSC genome and its functional annotations, we examined a 90-kb region on the distal X chromosome containing the *Myc* and *CG12535* loci (Figure 2B). For *Myc*, PRO-seq and Pol II ChIP-seq revealed robust transcription from a single promoter, likely regulated by four distinct enhancers identified by STARR-seq.

Direct RNA-seq reads revealed that *Myc* produces a single transcript isoform in OSCs with a 3′ UTR that is distinct from the annotated isoforms in FlyBase. This transcript is a source of abundant piRNAs that map almost exclusively to the 3′ UTR portion.

By contrast, *CG12535* showed no detectable promoter activity. Yet RNA-seq data revealed expression of its terminal two exons (Figure 2C). Closer inspection identified a heterozygous insertion of a *springer* LTR retrotransposon within the second intron as the source of this activity. The *springer* element is oriented in the same direction as the host gene and is not targeted by antisense piRNAs in OSCs. Genome-unique Pol II ChIP-seq reads together with PRO-seq data confirmed that the insertion is transcriptionally active. Moreover, direct RNA-seq demonstrated that transcription initiates at the first *springer* LTR and, through use of the *envelope* splice donor, splices into the last two *CG12535* exons to produce transposon-mRNA chimeric transcripts (similar chimeric transcripts have also been described in [37]).

Notably, the *CG12535* locus also contains a heterozygous insertion of the *envelope*-encoding *gypsy* retrovirus. In wild-type cells, this element is embedded within H3K9me2/3-marked heterochromatin and is transcriptionally silent, consistent with abundant *gypsy* antisense piRNAs in OSCs. Loss of Piwi-dependent heterochromatin not only derepressed the *gypsy* insertion (Pol II ChIP-seq) but also indirectly increased transcription from the neighboring *springer* element, resulting in higher levels of *springer*-*CG12535* chimeric transcripts (RNA-seq, direct RNA-seq) (Figure 2D).

Together, the OSC genome assembly and its integrated functional datasets establish a powerful framework to interrogate the interplay between transposons, gene regulation, and piRNA-mediated silencing, providing both a reference resource and a platform for new discoveries.

### The transposon landscape in OSCs and its regulation by the piRNA pathway

Transposable elements are mobile genetic elements that can copy and mobilize to new genomic sites, generating structural variation within the genome. A whole-genome comparison between the OSC_r1.01 assembly and the *Drosophila melanogaster* dm6 reference genome revealed more than 5,000 structural variants larger than 50 bp (Figure 3A). Most variants represented small insertions or deletions, but a prominent class, spanning 5-10 kb, consisted almost entirely of transposon insertions (Figure 3B). To systematically annotate these elements across the OSC genome, we applied RepeatMasker with a curated library of *D. melanogaster* transposon consensus sequences, identifying all transposon-derived genomic regions for subsequent analyses.

**Figure 3.**
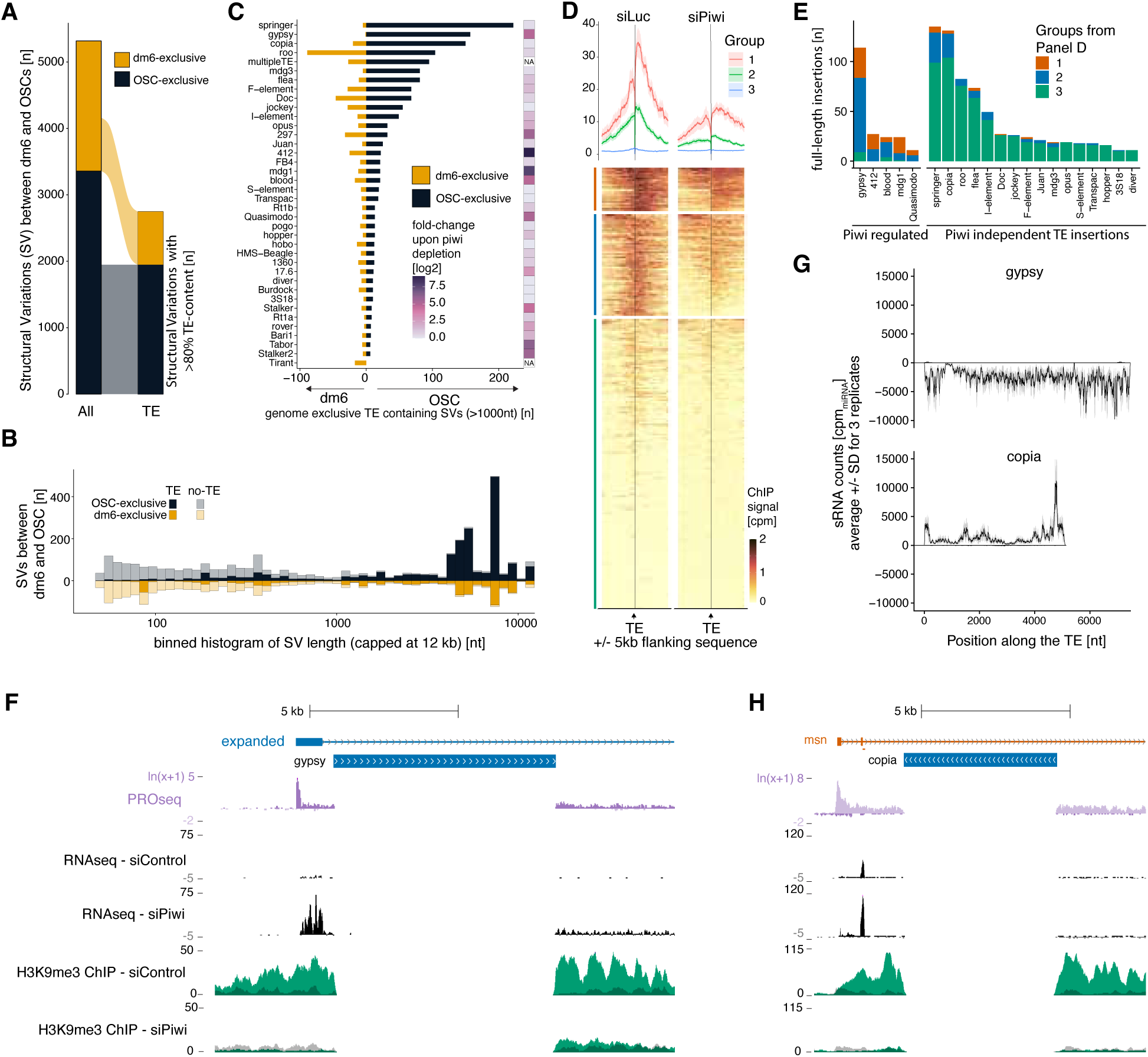
Transposon mobilization shapes the OSC genome and is influenced by piRNA-guided heterochromatin. (**A**) Quantification of structural variants (SVs) between the OSC and dm6 genomes. Data are presented for all SVs and for SVs comprising >80% transposable element content. Color grouping indicates the genome containing the additional sequence content. (**B**) Length distribution of the SVs as in A. SVs containing <80% transposable element content are shown shaded. (**C**) Structural variants (SVs) containing >80% transposable element content, shown per element for TEs with 10 or more variations. For each element, dm6-specific insertions are displayed on the left and OSC-specific insertions on the right. The accompanying heatmap indicates log₂ fold change in expression upon *piwi* depletion, quantified from three replicate RNA-seq experiments (siControl vs. siPiwi). (**D**) Heatmaps of H3K9me3 ChIP-seq signal ±5 kb around full-length (>90% of consensus length) transposon insertions in control (left) and *piwi*-depleted (right) conditions. Insertions were clustered into three groups by k-means, and the average ChIP-seq signal for each group is shown as a meta-profile above the heatmaps. (**E**) Per-transposon split of H3K9me3 grouping as in (D). Transposons were classified into *Piwi*-regulated or *Piwi*-independent categories based on a log₂ fold-change cutoff of >2.5, as in (C). (**F, H**) Genome browser view of the *expanded* (F) and *msn* (H) loci. (ChIP, PRO and RNA signal are shown as coverage per million reads - PRO-seq signal is displayed as ln(x + 1)-transformed coverage). ChIP-seq is shown as an overlay of ChIP signal (green) and ChIP input (grey). (**G**) Histogram plots showing piRNA coverage along the length of the *gypsy* and *copia* transposable elements. The x-axis indicates the position on the element, and the y-axis shows average piRNA counts from three replicates, normalized to 1 million miRNAs. Shaded areas represent the standard deviation across replicates.

Most of the transposon-generated structural variants represent novel insertions in the OSC genome absent in dm6. This is consistent with transposon mobilizations accompanying the immortalization of Drosophila cell lines [33] (Figure 3C, figure-supplement 3A). A few families disproportionately contributed to this process, including *springer*, *gypsy*, *copia*, *mdg3*, and *flea*. A striking example is the endogenous retrovirus *gypsy*: whereas the reference genome encodes only a single intact copy, the OSC genome contains more than 150 insertions. In contrast, *tirant*, an endogenous retrovirus that invaded natural *D. melanogaster* populations around the 1950s [38], is abundant in the reference genome but absent in OSCs, suggesting that OSCs were derived from *tirant*-naïve fly strains.

The OSC line expresses a functional nuclear Piwi/piRNA pathway, in which Piwi establishes repressive H3K9me2/3-marked heterochromatin domains by binding nascent transposon transcripts complementary to piRNAs [24]. With a comprehensive transposon insertion map in hand, we assessed the chromatin environment flanking all euchromatic insertions. Hierarchical clustering of H3K9me3 profiles in wildtype and Piwi-depleted cells grouped transposon insertions into three categories: (i) those associated with strong Piwi-dependent heterochromatin, (ii) those with weaker but still Piwi-dependent domains, and (iii) those with little or no heterochromatin in either condition (Figure 3D).

This grouping closely mirrored whether a transposon family is under piRNA control [24]. Families whose expression is strongly repressed by piRNAs in OSCs (e.g. *gypsy*, *mdg1*, *412*) were almost exclusively assigned to groups I and II (Figure 3E, F). By contrast, families either lacking piRNA targeting (e.g. *copia*, *springer*) or transcriptional activity in OSCs (e.g. *roo*, *Doc*, *mdg3*) were largely placed in group III, with no detectable heterochromatin. These results establish that the formation of Piwi-dependent heterochromatin requires both transcriptional activity of the transposon and the presence of antisense piRNAs against the element.

Unexpectedly, we also identified striking exceptions to this grouping within transposon families. One prominent example was a *copia* insertion within the *msn* gene, which formed the strongest Piwi-dependent heterochromatin domain in the genome. This was surprising because *copia* is highly expressed in OSCs and produces no antisense piRNAs consistent with most of its ∼150 genomic insertions lacking heterochromatin (Figure 3G). Closer inspection revealed that nearly all heterochromatin-marked *copia* insertions share a distinctive feature: they reside in antisense orientation within introns of actively transcribed host genes (figure-supplement 3B). In this context, the abundant sense *copia* piRNAs generated in OSCs can recognize nascent antisense *copia* transcripts and nucleate local heterochromatin domains within host gene introns (Figure 3H). We observed a similar pattern for *springer*, another highly expressed element in OSCs that generates abundant sense but hardly any antisense piRNAs.

Together, these results demonstrate how precise knowledge of transposon insertions, together with functional datasets, can aid the understanding of how the nuclear piRNA pathway impacts gene expression and chromatin patterns.

### The *flamenco* cluster directly determines transposon piRNA profiles in OSCs

The mechanisms that bias piRNA production toward antisense transposon sequences remain poorly understood. OSCs recapitulate the situation in the somatic follicular epithelium, where the *flamenco* locus has been identified as the principal genomic source of transposon antisense piRNAs [35, 39–42]. Located at the boundary between euchromatin and pericentromeric heterochromatin on the X chromosome, *flamenco* stood out in our comparison of the OSC genome assembly with the dm6 reference as one of the most structurally diverse regions (figure-supplement 4A), containing multiple medium-to large-scale sequence changes (figure-supplement 4B).

Alignment of Oxford Nanopore long reads obtained from OSC genomic DNA highlighted these differences. Whereas large portions of the ∼800 kb region between the *DIP1* and *CG14621* genes were fragmented and poorly covered in the alignments to dm6 (Figure 4A), reads aligned seamlessly to the OSC genome. Coverage was uniform, interrupted only by seven heterozygous structural variants, including a heterozygous *gypsy* insertion at the *flamenco* 5′ end (Figure 4B). The OSC genome also resolves the assembly gaps present in dm6, providing a contiguous reconstruction of the extended *flamenco* locus between *DIP1* and *CG14621*. Strikingly, this expands the general locus to a region of 1.5 Mb, raising the important question of how far functional *flamenco* transcription extends beyond its annotated promoter near *DIP1*.

**Figure 4.**
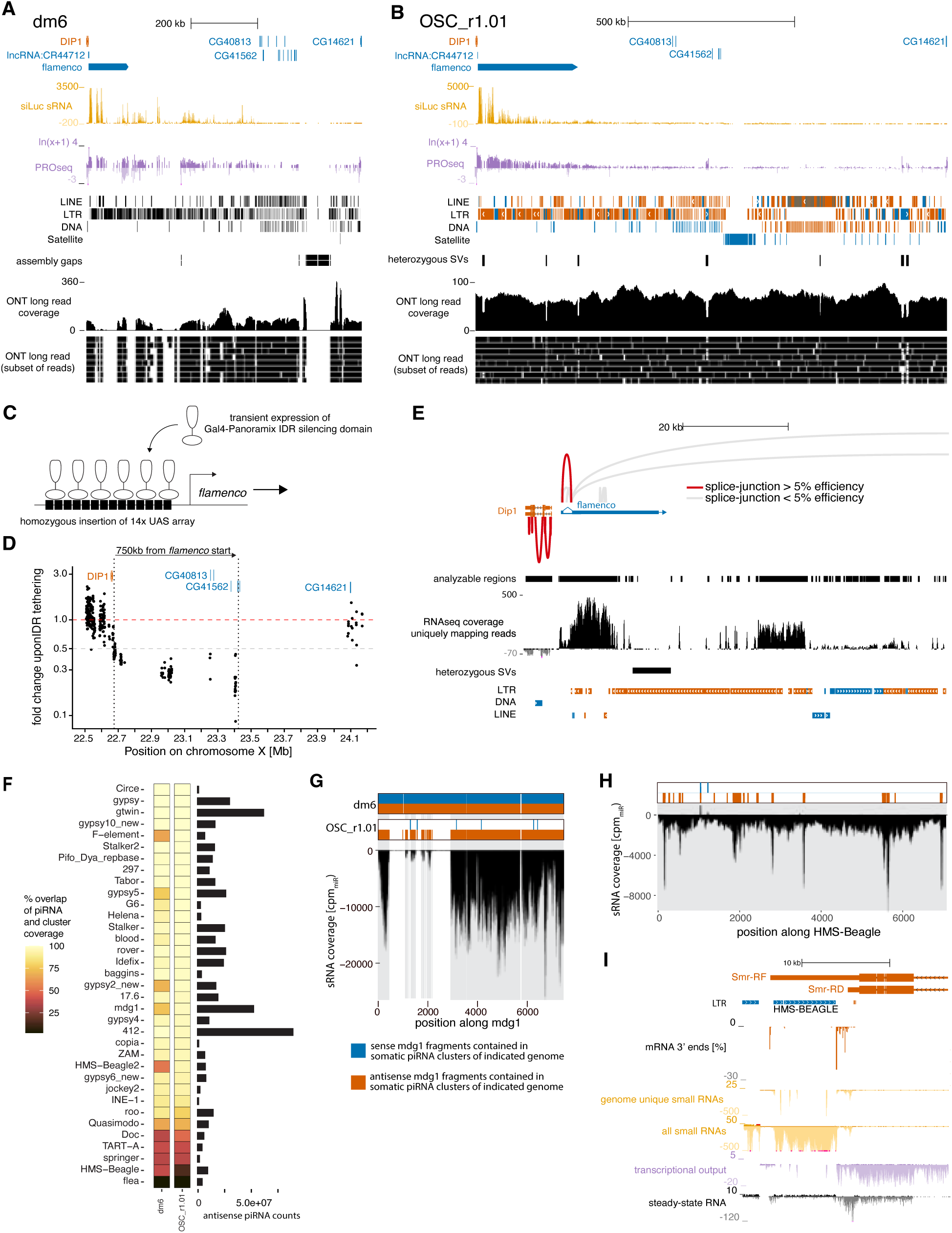
Reconstruction of the *flamenco* cluster in OSCs reveals its transcriptional extent and defines transposon piRNA landscapes (A) Genome browser view of the extended *flamenco* locus (*DIP1* to *CG14621*) in dm6. Small RNA data are normalized to 1 million miRNAs. PRO-seq signal is ln(x+1)-transformed coverage per million. ONT direct RNA-seq is shown as coverage and representative reads. Assembly gaps follow dm6 annotations. (B) Same region in the OSC genome. Small RNA normalized to 1 million miRNAs; PRO-seq as ln(x+1) per million. ONT direct RNA-seq shown as coverage and representative reads. Heterozygous structural variants from ONT genomic DNA (Sniffles) are indicated. (**C**) Schematic of the *flamenco* silencing experiment. (**D**) 1-kb tile analysis of fold change in small RNA upon IDR tethering at the *flamenco* promoter across the extended locus. Dotted lines mark the region from the *flamenco* TSS to the most distal analyzable point showing silencing. (**E**) Genome browser view of the initial 73 kb of *flamenco*, including *DIP1*. All identified splice junctions are shown. Analyzable (genome-unique) regions are indicated; multi-mapping regions were excluded from splice assessment. RNA-seq coverage used for splice analysis is shown as counts per million mapped reads. (**F**) Heatmap of concordance between antisense piRNA coverage of transposable elements (TEs) and their corresponding fragments within the *flamenco*, *20A*, and *77B* piRNA clusters, based on dm6 and OSC, respectively. Total piRNAs targeting each TE (per 1 million miRNAs) are shown alongside. (**G**) Histogram of piRNA coverage along *mdg1*. Above, *mdg1* fragments present in *flamenco*, *20A*, and *77B* (dm6 and OSC) are shown (sense: blue; antisense: orange). Regions with antisense piRNA coverage are shaded gray. (**H**) Histogram of piRNA coverage along *HMS-Beagle*. Above, genomic fragments present in *flamenco*, *20A*, and *77B* in the OSC genome are shown (sense: blue; antisense: orange). Regions with antisense piRNA coverage are shaded gray. (**I**) Genome browser view of the antisense *HMS-Beagle* insertion in the *Smr* 3′ UTR. Individual genic 3′ ends from ONT direct RNA-seq 3′-end quantification are shown as percent of all detected *Smr* 3′ ends. RNA-seq and PRO-seq are counts per million; small RNAs shown as genome-unique and all-mapping reads normalized to 1 million miRNAs.

To address this, we inserted UAS sites upstream of the *flamenco* promoter in both alleles of OSCs (Figure 4C). Expression of a fusion protein between the GAL4 DNA-binding domain and the transcriptional repressor domain of Panoramix selectively suppressed piRNA production from *flamenco* (Figure 4D) without affecting *cluster 20A*, *cluster 77B* (figure-supplement 4C, D), or piRNA-generating 3′ UTRs or elsewhere on the X chromosome (figure-supplement 4E, F, G). At *flamenco*, piRNA loss extended at least as far as ∼730 kb downstream of the promoter, demonstrating that transcription across much of the expanded *flamenco* locus contributes to piRNA biogenesis.

Closer inspection of uniquely mapping small RNAs revealed that piRNA output diminishes gradually along *flamenco*, dropping to below 5% beyond 500 kb from the transcription start site (Figure 4B). Because *flamenco* transcripts have been proposed to undergo extensive splicing [40], we considered whether alternative splicing could account for this decline. As direct RNA-seq was uninformative due to low coverage, we analyzed Illumina RNA-seq data. This confirmed the previously described intron near the *flamenco* promoter but provided no evidence for additional prevalent splice events across the locus, even in highly mappable regions (Figure 4E, figure supplement 4H). These findings argue that the drop in piRNA output reflects a gradual decay in transcriptional activity rather than alternative processing, a conclusion directly supported by PRO-seq data at the *flamenco* locus (Figure 4B).

We finally asked how the sequence content of the three active piRNA clusters in OSCs—*flamenco*, *20A*, and *77B*—shapes the transposon piRNA repertoire. Because clusters often contain fragmented transposon copies, we compared their transposon fragment content to the actual transposon piRNA profiles [10]. Using both dm6 and OSC genome assemblies, we quantified the fraction of each transposon’s sequence present in antisense orientation in the three piRNA clusters (Figure 4F). The OSC assembly markedly improved the correlation between cluster content and observed piRNA profiles. For example, the LTR retrotransposon *mdg1* generates an exclusively antisense piRNA profile with several pronounced gaps. In dm6, *flamenco* harbors complete sense and antisense *mdg1* copies, very much inconsistent with the piRNA data (Figure 4G). By contrast, the OSC genome contains only antisense fragments, which correspond precisely to regions of abundant antisense piRNA production.

Notably, the low abundance piRNAs mapping to *mdg1* nucleotides 1,000-2,100 matched small antisense fragments uniquely found in *cluster 20A*, which is transcribed at considerably lower levels than *flamenco* in OSCs.

Overall, the sequence content of the three OSC piRNA clusters accounts for the vast majority of transposon antisense piRNAs in OSCs. The few exceptions involved elements such as *flea* and *HMS-Beagle* (Figure 4H). While we could not identify a *flea* insertion in the OSC genome as possible piRNA source locus, an antisense *HMS-Beagle* insertion within the 3′ UTR of *Smr* likely accounts for most of the observed *HMS-Beagle* antisense piRNAs while the more prominent peaks in piRNA density are most likely explained by small antisense fragments in *flamenco* (Figure 4I).

Taken together, the OSC genome assembly enables a complete reconstruction of the *flamenco* cluster, reveals its transcriptional extent of at least 700 kb, and demonstrates how its sequence content defines piRNA output for most transposons. These findings support the central principle that in somatic ovarian cells, piRNA cluster content largely predetermines the transposon antisense piRNA repertoire, with additional contributions from transposon insertions in 3′ UTRs of expressed host genes.

## DISCUSSION

We present a high-quality *de novo* assembly of the *Drosophila* ovarian somatic cell genome, providing a reference for one of the most widely used cell systems in piRNA research. This resource closes an important gap between the biology of OSCs and reliance on the *Drosophila melanogaster* reference genome, which differs considerably in transposon content and piRNA cluster composition.

The OSC genome resolves the structure and transcriptional extent of the *flamenco* piRNA cluster, demonstrates that cluster sequence content directly determines the piRNA repertoire targeting transposons in this cell line, and clarifies how orientation and genomic context of transposon insertions dictate whether they are targeted by the nuclear Piwi pathway. These findings showcase the value of accurate genome assemblies for interpreting small RNA populations and their functional consequences.

One of the goals of this work was to establish the OSC genome assembly as a useful community platform. By providing comprehensive gene and transposon annotations together with an interactive UCSC genome browser hub, it enables seamless integration of diverse datasets. We anticipate that this accessibility will aid the design of targeted experiments, facilitate cross-study comparisons, and accelerate discovery in piRNA biology. We note that Siomi, Iwasaki and colleagues recently published a genome assembly of the OSC line used in their laboratory [37], which differs slightly from the line sequenced by us [33]. As their assembly lacks annotations and a browser-compatible resource, we hope that a common platform featuring both genome assemblies will be established in the future.

## MATERIALS & METHODS

### OSC cell culture

Ovarian somatic cells (OSCs) were maintained at 27 °C in a humidified incubator with 2.5% CO₂, as previously described [43, 44]. Cells were cultured in M3 basal medium supplemented with 10% fetal bovine serum (FBS), insulin, glutathione, and fly extract. For siRNA-mediated knockdown, OSCs were electroporated using either an Amaxa 2B nucleofector (Lonza) with Kit V and program T-029, or an Amaxa 4D nucleofector with Buffer SF and program DG150. A total of 4 × 10⁶ cells were transfected in 100 µL electroporation buffer supplemented with 4 µL siRNA (50 µM stock). A second transfection was performed 48 h after the initial transfection, and cells were harvested 96 h post the initial transfection for downstream analyses.

### Genomic DNA isolation

Genomic DNA was isolated from OSC cells using a modified protocol based on Jain et.al [45]. OSCs from T75 flasks were washed with PBS and lysed directly in 10 mL lysis buffer (200 mM NaCl, 100 mM Tris-HCl pH 8.5, 50 mM EDTA, 0.5% SDS, 50 µL Qiagen Puregene Proteinase K per 10 mL). The lysate was incubated at 60 °C for 5 h with occasional gentle mixing. After cooling, 100 µL Qiagen RNase A solution was added and incubated for 2 h at 37 °C. Following RNase treatment, the lysate was transferred to a Qiagen MaXtract high-density tube for sequential extractions. It was first mixed with 10 mL phenol (pH 8.0) by rotation (20 min, 20 rpm), then centrifuged (10 min, 1,500 g, 4 °C). The aqueous phase was transferred to a new MaXtract tube, mixed with 5 mL phenol (pH 8.0) and 5 mL chloroform, and rotated (10 min, 20 rpm) before a second centrifugation. The final aqueous phase was mixed with 7 mL isopropanol to precipitate DNA, which was then spooled with a sealed Pasteur pipette. The DNA was washed three times with 70% ethanol and resuspended in 100 µL Tris-HCl (pH 8.7) by overnight rotation at 4 °C for solubilization.

### Nanopore DNA library preparation and sequencing

Libraries were prepared using either the LSK108 ligation sequencing kit (Oxford Nanopore Technologies) or the RAD002 rapid sequencing kit. LSK108 libraries were constructed according to the manufacturer’s standard protocol to ensure efficient adapter ligation and clean-up. RAD002 libraries followed the modified protocol of Jain et al. [45]. In this approach, the amount of transposase used during the fragmentation step was reduced to minimize over-fragmentation and preserve long DNA fragments, while the incubation time with the adaptor-ligation mix was extended to improve adapter attachment efficiency. These modifications enabled the recovery of ultra-long DNA molecules (>100 kb). To further protect high–molecular weight DNA during handling, library clean-up conditions were optimized to reduce mechanical shearing, and all pipetting steps were performed with wide-bore tips using slow, minimal mixing.

Sequencing libraries were directly loaded onto R9.4 (FLO-MIN106) flow cells and ran on the MinION device, following the manufacturer’s guidelines for flow cell priming, sample loading, and temperature control. Sequencing performance was monitored in real time with MinKNOW, and runs were continued until active pore counts declined substantially.

### Nanopore RNA library preparation and sequencing

Total RNA was isolated from 90% confluent T75 flasks of OSC cells. Cells were washed three times with PBS, lysed with 10 ml of Trizol, and incubated on a shaker for 30–60 minutes. The lysate was transferred to a 15 ml Falcon tube containing phase lock gel, mixed with 2 ml of chloroform by careful inversion, and centrifuged at 4800 g for 10 minutes at 20°C using a swing-out rotor. The aqueous phase was transferred to a fresh Falcon tube, 7 ml of isopropanol was added, and the mixture was incubated at room temperature for 15 minutes. RNA was then pelleted by centrifugation at 20,000 g for 1 hour at 4°C in a fixed-angle rotor. The supernatant was removed, and the pellet was washed with 75% ethanol. After complete removal of ethanol, the RNA pellet was resuspended in 500 µL of nuclease-free water without pipetting for mixing and carefully transferred to an Eppendorf tube. RNA concentration was measured using Qubit.

mRNA was selected using oligo(dT) Dynabeads. A 200 µL aliquot of Dynabead slurry was washed twice with 2x binding buffer (10 mM Tris-HCl pH 7.5, 1.0 M LiCl, and 2 mM EDTA) and resuspended in 100 µL of 2x binding buffer. Total RNA was heat-denatured at 65°C for 5 minutes and snap-cooled on ice. Washed beads were added to the heat-denatured RNA, and the mixture was incubated with overhead rotation for 15 minutes. After incubation, the supernatant was removed, and the beads were washed three times with 500 µL of wash buffer (10 mM Tris-HCl pH 7.5, 0.15 M LiCl, and 1 mM EDTA), resuspending the beads each time. mRNA was eluted into 12 µL of nuclease-free water by incubation @ 80°C for 1 minute. mRNA concentration was measured using Qubit.

Nanopore direct RNA sequencing libraries were prepared from 10 µL of eluted mRNA following the RNA002 protocol (Oxford Nanopore Technologies). Half of the prepared library was used on a fresh flow cell for sequencing.

### Illumina DNA library preparation

Genomic DNA was fragmented using a Covaris E220 instrument with microTUBEs containing AFA fibers, following the manufacturer’s settings for a 400 bp peak size (Peak Incident Power = 140, Duty Factor = 10%, Cycles per Burst = 200, Treatment Time = 55 s). Size selection was performed with AMPure XP beads (Beckman Coulter). Long fragments were first removed by adding 53 µL of bead slurry per 100 µL of genomic DNA. Subsequently, an additional 25 µL of AMPure beads was added, and DNA fragments were purified according to the manufacturer’s protocol. Libraries were prepared using the NEBNext DNA Library Prep Kit (New England Biolabs) according to the manufacturer’s instructions, except for the final amplification step. For amplification, libraries were PCR-amplified using KAPA HiFi polymerase with EvaGreen fluorescence incorporated for real-time monitoring. PCR reactions were terminated once predetermined RFU thresholds were reached to prevent over-amplification. After purifications, samples were sequenced (PE100, Illumina HiSeq2000).

### Hi-C library preparation

Hi-C libraries were generated as previously described [46], with minor modifications.

OSCs were harvested, washed in PBS, and resuspended in M3 medium containing 1% formaldehyde at a density of 5 × 10⁶ cells/mL. After incubation for 10 min at room temperature (RT) with gentle rotation, 2 M Tris-HCl (pH 8.0) was added to a final concentration of 0.125 M to quench crosslinking. Samples were incubated for 5 min at RT, followed by 20 min on ice. Cells were pelleted, washed twice with PBS, and stored in aliquots of 5 × 10⁶ cells at –80 °C. Two OSC aliquots (5 × 10⁶ cells each; 1 × 10⁷ total) were processed in parallel. Cells were resuspended in 1 mL ice-cold lysis buffer, incubated for 30 min at 4 °C with rotation, and pelleted (2,500 g, 5 min, 4 °C). Pelleted nuclei were resuspended in 1 mL lysis buffer, pelleted again, and finally resuspended in 400 µL 1.2× NEB buffer 3.1, minimizing additional wash steps to reduce clumping.

Nuclei were digested overnight at 37 °C with rotation (800 rpm) using 30 U DpnII (NEB) in a final volume of 80 µL. After digestion, pellets were resuspended in 52 µL fill-in mix containing 2.2 µL biotin-14-dATP (1 mM, Jena Bioscience), 1.1 µL dNTP mix (2 mM dCTP, dGTP, dTTP each), 11 µL Klenow fragment, 5.72 µL NEB buffer 2, and nuclease-free water to volume, and incubated for 1.5 h at 37 °C with rotation. Filled-in nuclei were resuspended in a 250 µL ligation mix (27.5 µL Thermo T4 DNA ligase buffer, 2.2 µL 10% Triton X-100, 2.25 µL BSA [10 mg/mL], 232.5 µL H₂O, and 11 µL T4 DNA ligase).

Samples were incubated at RT for 5 h with rotation. Aliquots (10 µL) were collected at digestion and ligation steps as process controls. Post-ligation, reactions were diluted to 500 µL with PBS, supplemented with NaCl (500 mM final concentration) and 10 µL proteinase K (20 mg/mL), and incubated overnight at 65 °C. DNA was purified by two rounds of phenol–chloroform extraction, followed by a chloroform wash, and ethanol precipitation in the presence of glycogen. Pellets were resuspended in 150 µL H₂O (controls in 10 µL), treated with RNase A (2 µL [10 U/µL], 30 min, 37 °C), and quantified using Qubit. 130 microliters of DNA sample were transferred to Covaris microTUBEs and sheared on a Covaris E220 (<10 °C, intensifier inserted, 105 W peak power, 5% duty factor, 200 cycles/burst, 55 s). DNA was rinsed from Covaris tubes with 80 µL EB and subjected to two-step Ampure XP bead size selection (0.44× and 0.18×). DNA was eluted in 100 µL H₂O. Biotinylated DNA was captured with Dynabeads MyOne Streptavidin C1, pre-washed in Tween wash buffer (5 mM Tris-HCl pH 7.5, 0.5 mM EDTA, 1 M NaCl, 0.05% Tween-20). Beads were resuspended in 100 µL 2× binding buffer (10 mM Tris-HCl pH 7.5, 1 mM EDTA, 2 M NaCl), mixed with DNA, and incubated for 1.5 h at room temperature with rotation. Beads were washed twice in 500 µL Tween wash buffer, twice in H₂O, and resuspended in 50 µL H₂O. Bead-bound DNA was subjected to end repair with NEBNext Ultra II End Prep (7 µL End Prep buffer, 3 µL enzyme mix; 30 min at 20 °C then 30 min at 65 °C), adaptor ligation (2.5 µL NEB Illumina adaptor, 1 µL ligation enhancer, 30 µL ligation master mix; 15 min at 20 °C), and USER enzyme treatment (3 µL; 15 min, 37 °C). Beads were washed four times with 500 µL Tween wash buffer. DNA was eluted in 20 µL pre-warmed elution buffer (10 mM EDTA, 95% formamide, 65 °C), ethanol-precipitated with glycogen and NaCl, and resuspended in 15 µL H₂O. PCR amplification was performed with KAPA HiFi polymerase (25 µL master mix, 2 µL EvaGreen, 2.5 µL 10 µM barcoded primer, 1 µL 25 µM universal forward primer, 4.5 µL water). Cycling proceeded for 5 cycles, stopping when real-time fluorescence monitoring reached 3,600 RFU. PCR products were diluted, purified using Zymo DNA Clean & Concentrator-5, and eluted in 15 µL H₂O. Final libraries were quantified with Qubit, quality-checked on a Fragment Analyzer, and sequenced (PE75, illumina NextSeq 550).

### sRNAseq library preparation

sRNAseq libraries were prepared by first purifying Argonaute–sRNA complexes with TraPR resin columns [47]. OSCs were harvested using trypsin, washed 2x with PBS and stored at-70°C after snap freezing cell-pellets in liquid nitrogen. OSCs were lysed by adding 330 µL of TraPR lysis buffer. OSC lysates were clarified, normalized to equal protein amounts (depending on the experiment 70-120 µg total protein) after Bradford measurements, applied to TraPR columns, and Argonaute-bound RNAs were eluted, followed by phenol–chloroform extraction and precipitation. This column-based isolation bypasses conventional gel-based size selection and provides highly specific enrichment for Ago-associated small RNAs. To minimize ligation bias and facilitate library multiplexing, adaptors were designed with random nucleotides placed adjacent to the ligating end [48]. The 3′ adaptor carried six random bases at its 5′ terminus and contained sample barcodes (X) for multiplexing (5rApp/NNNNNNXXXX–Adaptor–/3ddC). The 5′ adaptor was an RNA oligo bearing four random nucleotides at its 3′ terminus (…UCUNNNN). After 3′ adaptor ligation with T4 RNA ligase 2, truncated KQ, samples were pooled as necessary, spiked with fluorescently labeled and ligation-blocked oligos as size controls, and gel-purified on a 12.5% polyacrylamide–urea gel. Small RNAs were isolated using the Zymo ZR small RNA PAGE Recovery kit and subjected to 5′ adaptor ligation with T4 RNA ligase 1. Following cleanup with Zymo RCC5 columns, reverse transcription was performed. Libraries were PCR-amplified with KAPA HiFi polymerase and dual-index barcoded primers. Amplification was monitored in real time with EvaGreen, and reactions were stopped upon reaching 3,500–4,000 RFU to prevent overamplification. Final libraries were size-selected on low-melt agarose gels, gel-purified (Zymo), quantified with Qubit, multiplexed and sequenced (SE50, Illumina NovaSeq6000).

### PROseq library preparation

PROseq libraries were prepared using a protocol modified from Mahat et al. [49] from both isolated nuclei and permeabilized cells. Cells were harvested, washed with ice-cold PBS, and counted. Nuclei were isolated by resuspension in douncing buffer (10 mM Tris-HCl pH 7.4, 300 mM Sucrose, 3 mM CaCl₂, 2 mM MgCl₂, 0.1% Triton X-100, supplemented with 0.5 mM DTT, protease and RNase inhibitors), dounced 25 times with a tight pestle, and pelleted. Permeabilized cells were prepared by resuspension in permeabilization buffer (10 mM Tris-HCl pH 7.4, 300 mM Sucrose, 10 mM KCl, 5 mM MgCl₂, 1 mM EGTA, 0.05% Tween, 0.1% NP40 substitute, supplemented with 0.5 mM DTT, protease, and RNase inhibitors), incubated on ice for 5 minutes, and pelleted.

Both nuclei and permeabilized cells were resuspended in storage buffer (10 mM Tris-HCl pH 8.0, 25% glycerol, 5 mM MgCl₂, 0.1 mM EDTA, supplemented with 5 mM DTT), aliquoted, and snap-frozen in liquid nitrogen. Nuclear run-on reactions were performed using 2xNRO buffer supplemented with biotin-11-CTP and biotin-11-UTP (for 2-biotin mixes) or single biotinylated nucleotides (for 1-biotin mixes), ATP, GTP, and RNaseOUT. Samples were incubated at 30°C for 3 minutes, followed by RNA extraction using Trizol-LS and chloroform. RNA was precipitated with Glycoblue and 100% ethanol, incubated for 15 minutes at room temperature, and pelleted by centrifugation (16000 x G, 1 hour, 4°C). Pellets were washed with 75% ethanol and stored at-80°C. Isolated RNA was fragmented by heat denaturation (65°C for 40 seconds) and alkaline hydrolysis (1N NaOH for 10 minutes on ice), then purified using P30 columns.

Biotinylated RNA was enriched using MyOne C1 streptavidin beads, washed with high-salt, PROseq binding, and low-salt buffers, and eluted with Trizol and chloroform extraction. RNA was precipitated with Glycoblue and 100% ethanol. After decapping and hydroxyl repair using TAP and PNK enzymes followed by phenol purification, libraries were generated under sRNA-seq ligation conditions. 3′ adaptor ligation was performed with T4 RNA ligase 2, truncated KQ, using adaptors containing four random nucleotides at the 5′ end [48]. After a second streptavidin purification, 5′ adaptor ligation was performed with T4 RNA ligase, using adaptors containing four random nucleotides at the 3′ end, followed by a third streptavidin purification. Reverse transcription was carried out with SuperScript II using an adaptor-specific primer. Libraries were PCR-amplified with KAPA HiFi polymerase and barcoded Illumina primers in reactions monitored by EvaGreen real-time fluorescence. Reactions were stopped after reaching defined RFU thresholds (12–17 cycles). Products were separated on 2.5% agarose gels, gel-purified (Zymo), and multiplexed for sequencing (PE50, Illumina HiSeq2500).

### RNAseq library preparation

Total RNA was isolated using Trizol (Invitrogen) according to the manufacturer’s instructions. Ribosomal RNA was depleted using an RNAseH based protocol [50, 51]. In brief, standard 50-mer DNA oligos complementary to rRNA precursor and mitochondrial rRNA sequences were annealed to 1 µg total RNA in Hybridase buffer with EDTA. Samples were denatured (95 °C, 3 min), cooled gradually to 45 °C, and incubated with Thermostable Hybridase RNAseH (1 h, 45 °C) to specifically degrade rRNA. Following digestion, samples were treated with Turbo DNAse (30 min, 37 °C) and purified using RNA Clean & Concentrator-5 columns (Zymo). Elution was carried out in NEBNext Ultra II First Strand Synthesis buffer for downstream library preparation. RNA-seq libraries were generated using the NEBNext Ultra II Directional RNA Library Prep Kit (New England Biolabs) according to the manufacturer’s instructions, except for the final amplification. Libraries were PCR-amplified with KAPA HiFi polymerase and barcoded Illumina primers, with amplification monitored in real time using EvaGreen fluorescence. Reactions were stopped once predetermined RFU thresholds were reached to avoid over-amplification. Libraries were then purified, quantified by Qubit, assessed for fragment size, and multiplexed for Illumina sequencing.

### Generation of UAS landing-site cell lines

A donor repair template was generated by cloning ∼1 kb homology arms flanking an attP-flanked Puro-mCherry cassette into a plasmid backbone. The construct was assembled by Gibson assembly, propagated in *E. coli* STBL3, and sequence-verified by Sanger sequencing. A CRISPR-Cas9 sgRNA targeting the chosen insertion site within the *flamenco* upstream region (target: TATAAAAGTTACAAAATACG) was designed using CHOPCHOP, synthesized as complementary oligonucleotides, and cloned into a Cas9 expression vector (Addgene 49330 [52]). OSCs were co-transfected with the verified donor and sgRNA/Cas9 plasmids using the Amaxa 4D nucleofection system (buffer SF, program DG150). Forty-eight hours post-transfection, cells were replated into 10-cm dishes at varying densities (2.5%, 5%, 10% of total cells). Selection was initiated with puromycin (1:1500) five days after plating. Resistant colonies were picked after 7 days, expanded in 96-well plates, and subsequently scaled up. Clonal integration was assessed by long-range PCR spanning the *integration site* and ddPCR using primers specific for the donor cassette and endogenous controls. The assays confirmed both the integrity and zygosity of the engineered landing site.

### *flamenco* silencing

A plasmid containing an attB–selection cassette–14xUAS–attB was constructed. The selection cassette encoded *mCherry–Blasticidin*, driven by the *D. yakuba tj* enhancer with the *armi* 3′ UTR. Homozygous *flamenco* landing site (LS) OSCs were co-transfected with the UAS construct and a φC31 integrase expression plasmid using Amaxa 4D nucleofection (buffer SF, program DG150). Five days post-transfection, feeder OSCs were prepared by diluting wild-type OSCs to 1 × 10^6^ cells/mL, irradiating twice for 20 min with mixing between treatments, and plating 1 × 10^5^ feeder cells per well of a 96-well plate. The following day, single cells that were GFP positive and mCherry negative were sorted into feeder-containing wells. After 14 days, wells with visible colonies were identified by plate scanning (Sapphire - Azure Biosystems) and expanded. Genomic PCR was used to assess cassette integration and zygosity, and Sanger sequencing confirmed integration orientation. Clones homozygous for UAS insertions oriented towards *flamenco* were subsequently transfected with tethering plasmids encoding either Gal4 alone or Gal4 fused to the Panoramix IDR silencing domain [53]. Transfections were repeated after 48h to sustain tethering. At 6 and 7 days post transfection, cells were harvested by FACS, gated on reduced GFP reporter signal from the co-silenced selection cassette. For Gal4-only controls, no FACS enrichment was applied. Small RNAs were isolated from sorted cells and sequenced as described in the small RNA sequencing section, except that the 3′ adapter lacked a barcode and carried only four random nucleotides at its 5′ end.

### Nanopore DNA sequencing base-calling and read processing

Raw multi-read FAST5 files were base-called on GPUs with ONT Guppy v5.0.7+2332e8d65 using the super-accuracy model dna_r9.4.1_450bps_sup (Oxford Nanopore Technologies). Base-calling was performed without quality filtering to retain all reads for downstream QC and filtering (--disable_qscore_filtering). Post-base-calling, FASTQ files from each batch (including pass and fail subdirectories) were concatenated and processed in parallel to generate adapter-trimmed and untrimmed read sets.

Adapter trimming was performed with Porechop [54] using default settings with minimum split read size of 1,000 nt and adapter detection on a 1,000-read subsample (--min_split_read_size 1000 --check_reads 1000). For the untrimmed set, chimeric reads were discarded (--discard_middle) but end adapters were retained. Read length and average per-base quality scores were computed from FASTQ quality strings using a precomputed Phred score lookup table, and appended to read headers (LENGTH=, avgQUAL=) with seqkit [55] and mawk.

### Nanopore direct RNA sequencing base-calling and read processing

Direct RNA multi-read FAST5 files were base-called on GPUs with ONT Guppy v5.0.7+2332e8d65 using the appropriate direct RNA configuration for the flow cell/kit (Oxford Nanopore Technologies). Base calling emitted all reads (-- disable_qscore_filtering) while preserving raw signal and disabling base-caller trimming where applicable (--fast5_out --trim_strategy none). Post-base-calling, per-batch FASTQ files were concatenated and processed to annotate read bodies upstream of the poly(A) tail and to estimate poly(A) lengths. Base-called FAST5 files were converted to gzip-compressed FAST5 (compress_fast5, ONT) and poly(A) tails were identified with tailfindr in R, yielding per-read tail metrics (tail_end_base_index, tail_length) [56]. For each read, sequence and quality strings were truncated at tail_end_base_index to retain the transcribed body upstream of the poly(A) tract, and metadata were appended to read headers (LENGTH=, avgQUAL=, polyAlength=) with seqkit [55] and mawk.

Reads without a detectable tail were flagged (polyAlength=notDetected) and listed for QC. For 3′ UTR annotation, only direct RNA reads with a detected poly(A) tail were used.

### Genome assembly and haplotig purging

Nanopore reads were filtered to retain sequences >15,000 nt, scrubbed with Yacrd v0.6.2 (-c 4-n 0.4 scrubb) [57], and quality-trimmed with Filtlong v0.2.0 (--target_bases 14400000000; https://github.com/rrwick/Filtlong) to obtain approximately 100× coverage. Filtered reads were assembled with Shasta @ 3367f960 using custom settings (see associated code deposition)[58]. Redundant haplotigs were identified and removed with Purge Haplotigs [59] (modified version https://gitlab.com/dominik-handler/purgehaplotigs@2b6eb0f2). Read-depth histograms were generated by mapping reads with Minimap2 v2.18 [60] and processed with purge_haplotigs hist.

Repeat-masked regions were annotated with RepeatMasker v4.1.0 (Smit, AFA, Hubley, R & Green, P. *RepeatMasker Open-4.0*.2013-2015 http://www.repeatmasker.org) (NCBI/RMBLAST v2.10.0+, CONS-Dfam_3.1-rb2018102 [61]). Haplotig purging was performed using purge_haplotigs cov-l 10-m 48-h 180, followed by purging with repeat annotations taken into account. Overlapping contig ends were trimmed with purge_haplotigs clip using default settings. Missasembly correction was performed based on read-alignment of Oxford Nanopore Technologies (ONT) reads and analysis of read-coverage gaps. ONT long reads were aligned to the initial assembly using minimap2 (map-ont mode) with secondary alignments excluded. Alignments were filtered to remove unmapped and supplementary reads, converted to BED format using samtools [62] and bedtools [63], and sorted by genomic coordinates. Coverage gaps indicative of potential mis-assemblies were identified using bedtools genomecov to generate per-base coverage across all contigs. Regions with coverage below 10× and located more than 30 kb from contig ends were flagged as candidate mis-assembly sites, provided the gap spanned at least 20 bp. Adjacent gaps within 1 kb were merged using bedtools merge to define consolidated break regions. For contigs containing identified gaps, splitting coordinates were determined by creating intervals between consecutive gap regions, with each resulting fragment assigned a unique identifier.

Contigs without detected mis-assemblies were retained intact with their original identifiers. The final split coordinates were used with bedtools getfasta to extract individual contig fragments, generating a corrected assembly file for subsequent Hi-C scaffolding. After mis-assembly correction, scaffolding was performed using Hi-C contact data. Raw Hi-C read pairs were aligned to the split genome assembly with BWA-MEM [64] using parameters optimized for Hi-C data (-A 1-B 4-E 50-L 0).

Alignments were filtered to remove low-quality reads and artifacts with the filter_five_end.pl script provided in the SALSA2 package [65, 66] then combined and sorted [65, 66]. Read groups were assigned with Picard AddOrReplaceReadGroups (“Picard Toolkit.” 2019. Broad Institute, GitHub

Repository. https://broadinstitute.github.io/picard/; Broad Institute), and PCR duplicates were identified and removed using Picard MarkDuplicates. Scaffolding was then carried out with SALSA2, specifying GATC as the restriction enzyme motif to match the DpnII digestion used during Hi-C library preparation. Additional super-scaffolding was performed against the *Drosophila melanogaster* dm6 reference genome using RagTag (v2.1.0) [67] with default parameters. Gap-filling was then carried out in two successive rounds using an in-house pipeline supported by Oxford Nanopore long reads. Assembly gaps were first identified and flanking regions (20 kb on each side) extracted. Long reads were mapped to the flanks with minimap2 (map-ont mode), and candidate gap-spanning reads were selected. When a single bridging read was available, it was polished with racon to generate a gap-filling contig. In the absence of bridging reads, flanking reads were assembled de novo using wtdbg2/WTPOA [68] to produce contigs spanning both flanks. Candidate contigs were validated by realignment of the flanking regions and, if consistent, used to replace the corresponding gap sequence.

Transposable element (TE) sequences were systematically integrated into the reference genome assembly using a custom computational pipeline. Nanopore reads were mapped to the reference assembly using minimap2 with Oxford Nanopore Technologies (ONT) parameters. Structural variants, particularly large insertions (>100 bp), were identified using Sniffles [69, 70] and phased to distinguish allelic variants. TE-containing reads were identified by mapping reads against a TE consensus library using minimap2, with flanking regions (default 1 kb) extracted to distinguish between TEs already present in the genome versus those absent from the assembly. Flanking sequences extracted from TE-containing reads were mapped back to the reference genome to classify TEs as either genome present or absent. TEs were categorized based on distance between flanking region mappings: those with gaps <50 bp were classified as absent, those with gaps matching expected TE size (±20%) were classified as present, and others flagged for manual review. Coverage analysis identified genomic regions with high-confidence missing TEs using a minimum coverage threshold of 4 supporting reads. Regions were merged within 20 bp windows and coordinates adjusted for subsequent insertions.

Phase information from the pre-phased structural variants was utilized to ensure proper allelic assignment of TE insertions. For each identified missing TE locus, relevant ONT read segments were extracted and assembled using wtdbg2 followed by consensus polishing with racon [71]. Assembled contigs containing both upstream and downstream flanking sequences were identified and validated against the genome. Successfully assembled TEs with valid flanking sequences were integrated into the genome assembly by replacing the gap region with the assembled TE sequence.

Polishing of draft assemblies was performed in three successive stages using both Oxford Nanopore Technologies (ONT) long reads and Illumina short reads. First, Clair3 polishing with ONT reads [72]. ONT reads were aligned to the draft assembly with minimap2 v2.24 using the map-ont preset, and alignments were sorted and indexed with samtools v1.14. Variants were called with Clair3 v0.1-r12 using the ONT model and the --enable_long_indel option. Homozygous alternate variants (1/1) flagged as PASS with insertion or deletion lengths <350 bp were retained. Filtered variants were applied with bcftools consensus v1.14 [62] to produce the Clair3-polished assembly. The Clair3-polished assembly was then further polished using the PEPPER-Margin-DeepVariant pipeline [73]. ONT reads were re-aligned to the Clair3-polished assembly with minimap2, and variants were called using the ONT R9.4.1 Guppy5 SUP model.

Homozygous alternate variants were filtered and applied with bcftools consensus, generating the PEPPER-DeepVariant–polished assembly. For the third round of polishing, Illumina paired-end reads were aligned to the Pepper-Margin-DeepVariant polished assembly with bwa-mem, and duplicates were marked with samtools. Variants were called with DeepVariant v1.1.0 [74] in whole-genome mode, filtered for homozygous alternate alleles, and applied with bcftools consensus to generate the final Illumina-DeepVariant–polished assembly.

### Genome annotation / UCSC hub generation

To annotate the assembled genome and generate a browser hub for visualization in the UCSC Genome Browser [75], we implemented a custom multi-step pipeline integrating gene annotation, repeat masking, read mapping, uniqueness profiling, and variant detection.

#### Assembly Preprocessing and Hub Initialization

Genome assemblies were preprocessed to ensure compatibility with downstream tools. Sequence identifiers containing special characters were sanitized using seqkit v2.3.0, and chromosome size files were generated. Assemblies were converted to 2bit format using faToTwoBit (UCSC Kent utilities [36, 75]) to support hub configuration.

#### Gene Annotation Strategy

A multi-tiered annotation strategy combined homology-based transfer, transcript alignment, and de novo gene prediction:

- Lift-over annotation: Reference annotations from *Drosophila melanogaster* (dm6) [76, 77] were transferred with Liftoff v1.6.3 [78], which uses sequence alignment to identify homologous loci and project gene models, while resolving structural variations. Pseudogenes were reclassified as mRNA features for consistency in downstream processing.
- Transcript mapping: FlyBase[77] transcript sequences (mRNA, CDS, UTRs, tRNAs, miRNAs, ncRNAs, pseudogenes) were re-aligned to the assembly. Y-linked transcripts were excluded unless explicitly requested.

○ Full-length transcripts and CDS were aligned using BLAT (parameters: - stepSize=5-fine-q=dna) [79], partitioned for parallel execution. Alignments covering ≥90% of transcript length were retained.
○ Short elements (<50 bp), such as UTRs, were aligned using STAR [80] with a genome index built from the target assembly.
○ BLAT and STAR results were merged, correcting spurious micro-introns (<20 bp) and enforcing accurate CDS boundaries.
○ The final transcript set was exported in bigGenePred format, validated with *genePredCheck*, and indexed (ixIxx) for fast UCSC search integration.
- De novo gene prediction: Long-read ONT RNA-seq and Illumina short-read RNA-seq data were used for ab initio discovery.

○ ONT RNA reads were aligned with minimap2 (splice-aware mode), and Illumina reads with HISAT2 v2.2.1 [81].
○ Transcripts were assembled with StringTie v2.2.1 [82] using mixed-mode analysis (minimum anchor length = 4 bp, minimum junction coverage = 2 reads, minimum transcript coverage = 3.9 reads per base).
- Annotation integration: Gene models from lift-over, transcript mapping, and de novo prediction were merged with a custom algorithm that prioritized evidence-supported annotations and resolved overlaps.

#### Repeat Element Annotation

Repetitive elements were annotated using RepeatMasker v4.1.2 with the Dfam database [61] plus a custom transposable element (TE) library. Outputs were converted into bigBed tracks for efficient UCSC visualization.

#### Read Mapping and Coverage Tracks (ONT)

Raw ONT DNA sequences were aligned using minimap2 (-ax map-ont), while ONT direct RNA and cDNA reads were mapped with splice-aware parameters (-ax splice-uf - k14). Alignments were sorted with samtools, and coverage tracks were generated using mosdepth. Normalized depth profiles were converted into bigWig format for direct genome browser display.

#### Genome Uniqueness Mapping

Uniqueness profiles were generated by sliding-window k-mer analysis. K-mers (25–50 nt) from both strands were generated using seqkit sliding. Single-copy k-mers were remapped to the assembly with Bowtie v1.3.1 [83] under multiple mismatch thresholds (0–3). The 5′ positions of uniquely mapped k-mers were extracted, converted to bigBed format, and organized into UCSC tracks representing regions of unique mappability.

#### Variant Detection

- ONT-based variants: ONT DNA reads were aligned with minimap2 (-Lax map-ont--secondary=no). SNVs and indels were identified with Clair3 [72] and PEPPER-DeepVariant [73], while structural variants were detected with Sniffles v2 [69, 70] and cuteSV [84]. Variants were processed with *bgzip* and *tabix*, stratified by zygosity, and summarized with rtg vcfstats (https://github.com/RealTimeGenomics/rtg-tools). Outputs were integrated as vcfTabix tracks.
- Illumina-based variants: Illumina reads were mapped with BWA-MEM[64] followed by filtering (samtools view-F 256 [62]), mate-pair correction (*samtools fixmate*), coordinate sorting (*samtools sort*), and duplicate marking (*samtools markdup*). Variants were called using DeepVariant [74] (WGS model), processed identically to ONT calls, and visualized in the UCSC hub.

#### UCSC Hub Construction

All annotation and analysis outputs (gene models, repeats, coverage, uniqueness, and variants) were converted into UCSC-compatible formats (bigWig, bigBed, bigGenePred, vcfTabix). Hub configuration files (*hub.txt, genomes.txt, trackDb.txt*) were automatically generated with metadata, methodology notes, and visualization defaults. Tracks were grouped and hierarchically organized to facilitate comparative exploration of genome structure, expression, and variation.

### 3’ UTR isoform refinement using ONT direct RNAseq reads

We developed a computational pipeline to refine 3′ UTR annotations and quantify 3′ UTR isoform usage by leveraging 3′ termini from Oxford Nanopore Technologies (ONT) direct RNA-seq reads. Our approach involved five key steps: (i) anchoring 3′ UTRs to coding sequence (CDS) stop codons, (ii) defining a conservative search window per transcript to exclude downstream genic and repetitive interference, (iii) identifying candidate 3′ ends from ONT read termini, (iv) tiling and quantifying local 3′ end signal, and (v) applying sequential filters to remove artifacts and low-confidence sites, ultimately yielding high-confidence 3′ UTR isoforms. Transcriptional stop codon positions, derived from expressed gene models, served as the initial anchors for 3′ UTR definition. Subsequently, UTR search intervals were established downstream of stop codons for sense strands and upstream for antisense strands. These windows were refined by considering the longest observed UTRs per gene and by intersecting with expressed annotations and repeat elements to prevent interference from unrelated genomic features. To identify candidate 3′ ends, ONT direct RNA-seq read 3′ termini were extracted from alignments. These positions were then grouped into “tiles” based on local read support, and each tile was quantified for total reads, peak reads, tile-specific reads, and fractional contribution.

For primary filtering, two strategies were employed:

- Continuous “RAMP-down” selection: This method iteratively retained tiles with significant read support, prioritizing pronounced drops in abundance consistent with major 3′ ends.
- Fraction-based filtering: Tiles were retained if they met predefined minimum read count thresholds and either a minimum fractional contribution or a minimum absolute read count.

Finally, to reduce redundancy, closely spaced minor peaks were either merged or discarded if they were within a defined proximity of a more abundant neighbor.

### Genome statistic comparison

Assembly quality and contiguity (Nx statistics) were assessed with QUAST [85]. Both the *Drosophila melanogaster* reference genome (dm6) and the OSC_r1.01 assembly were analyzed using the FlyBase reference annotation as a feature set. QUAST was run in large-genome mode with scaffold splitting, conserved gene identification, and eukaryote-specific settings (--large --features flybase.gff --split-scaffolds --conserved-genes-finding --eukaryote).

### BUSCO

Assembly completeness was evaluated using BUSCO v5 [86] with the *drosophila_odb12* lineage dataset in genome mode. BUSCO summary statistics and completeness plots were generated from the resulting output files.

### Whole-genome alignment and synteny visualization

Pairwise whole-genome alignments between each target assembly and the *Drosophila melanogaster* reference genome (dm6) were performed using NUCmer [87] with the parameters --maxmatch-c 100-b 500-l 50, which allow for sensitive detection of maximal exact matches and robust alignment of complex regions. Filtered alignments were converted to PAF format using *paftools delta2paf* [88] to enable downstream analysis and visualization. Repetitive versus unique alignments were classified by calculating mapping multiplicity of both query and reference intervals: alignments with multiple mappings were annotated as “multi”, whereas those mapping uniquely were retained as “uniq”. Dot plots were generated using a modified version of *paf2dotplot* (https://github.com/moold/paf2dotplot) that incorporated repeat annotations for color-coding and a custom color-scheme. Genome-wide synteny plots were created under standard filtering thresholds (minimum alignment length of 5 kb; minimum reference span of 100 kb). For targeted locus analysis, higher-resolution dot plots (minimum alignment length 1 kb; minimum reference span 1 kb) were produced, focusing on the extended flamenco region on chromosome X. For detailed analysis of the *flamenco* piRNA cluster, locus-specific sequences were extracted from dm6 and OSC_r1.01.

Pairwise NUCmer alignments were performed, filtered with delta-filter, and analyzed for structural rearrangements using SyRI (--nosnp --tdgaplen 10000) [89]. Local synteny plots were generated with plotsr (-s 5) [90] to visualize rearrangements within the *flamenco* cluster. For detailed analysis of the *flamenco* piRNA cluster, locus-specific sequences were extracted from dm6 and OSC_r1.01. Pairwise NUCmer alignments were performed, filtered with delta-filter, and analyzed for structural rearrangements using SyRI (--nosnp --tdgaplen 10000) [89]. Local synteny plots were generated with plotsr (-s 5) [90] to visualize rearrangements within the *flamenco* cluster.

### Hi-C data analysis

Hi-C sequencing reads were mapped to the OSC_r1.01 genome assembly using BWA-MEM [64] with the-P option for Hi-C paired-end data. Alignments were processed using samtools [62] and parsed with pairtools (v.1.1.3) [91]. Duplicate fragments were filtered with pairtools dedup to generate sets of valid pairs, duplicates, unmapped reads, and summary statistics. Hi-C sequencing reads were mapped to the OSC_r1.01 genome assembly using BWA-MEM [64] with the-P option for Hi-C paired-end data. Alignments were processed using samtools [62] and parsed with pairtools (v.1.1.3) [91]. Duplicate fragments were filtered with pairtools dedup to generate sets of valid pairs, duplicates, unmapped reads, and summary statistics. For chromosome-scale contact map construction, valid Hi-C pairs were converted into a balanced matrix format using Cooler (v.0.10.4) [92]. Bin intervals were generated at 1 kb resolution with cooler makebins.

Matrices were built with cooler cload pairs and balanced using cooler balance with a maximum median absolute deviation (MAD) threshold of 5. To enable multi-resolution analysis of genome architecture, balanced matrices were zoomified with cooler zoomify at 1 kb, 2 kb, 5 kb, 10 kb, 25 kb, 50 kb, and 100 kb resolutions. The resulting.mcool files were used for downstream visualization and quantitative analysis of Hi-C contact patterns. Hi-C contact matrix was visualized using HiGlass [93]. For chromosome-scale contact map construction, valid Hi-C pairs were converted into a balanced matrix format using Cooler (v.0.10.4) [92]. Bin intervals were generated at 1 kb resolution with cooler makebins. Matrices were built with cooler cload pairs and balanced using cooler balance with a maximum median absolute deviation (MAD) threshold of 5. To enable multi-resolution analysis of genome architecture, balanced matrices were zoomified with cooler zoomify at 1 kb, 2 kb, 5 kb, 10 kb, 25 kb, 50 kb, and 100 kb resolutions. The resulting.mcool files were used for downstream visualization and quantitative analysis of Hi-C contact patterns. Hi-C contact matrix was visualized using HiGlass [93].

### SNV detection

Illumina DNA-seq reads were aligned to the genomes using BWA-MEM [64], followed by sorting, indexing, and filtering for uniquely mapped reads with samtools (alignments with MAPQ <1 or alternative mappings were excluded) [62]. Variant calling was performed using DeepVariant (WGS model) [74] with assembly-specific references to generate VCF outputs and summarized with rtg vcfstats. Illumina DNA-seq reads were aligned to the genomes using BWA-MEM [64], followed by sorting, indexing, and filtering for uniquely mapped reads with samtools (alignments with MAPQ <1 or alternative mappings were excluded) [62]. Variant calling was performed using DeepVariant (WGS model) [74] with assembly-specific references to generate VCF outputs and summarized with rtg vcfstats. To quantify local SNV density, assemblies were partitioned into 1-kb windows using bedtools and homozygous and heterozygous SNV sets were separated using bcftools. Genome-uniqueness-filtered (>50% genome unique) windows were intersected with homozygous and heterozygous SNVs to obtain per-window counts, enabling assessment of genome-wide and locus-specific variant distributions.

### SV detection

Oxford Nanopore DNA reads were aligned to assemblies using minimap2 (-Lax map-ont -- secondary=no) and sorted/indexed with samtools. Structural variants (SVs) were identified using Sniffles v2 [69, 70] with default settings and summarized with rtg vcfstats. To classify SVs as transposable element (TE)-associated or not, insertion and deletion sequences >50 bp were extracted from VCF entries and aligned against a curated TE consensus library using minimap2. SVs with ≥80% sequence coverage from TE alignments were annotated as TE-related.

### Inter-genome SV detection

Structural variants (SVs) were identified by comparing the OSC genome assembly to the reference dm6 genome using minimap2 (v2.24) with parameters optimized for assembly-to-assembly alignment (-a-x asm5--cs-r2k). Alignments were sorted and indexed using samtools (v1.15). SVs were called using SVIM-asm [94] in haploid mode. To determine the zygosity of identified SVs in the OSC cell line, Oxford Nanopore Technologies (ONT) long-read sequencing data was mapped to the OSC assembly using minimap2 with ONT-specific parameters (-Lax map-ont --secondary=no).

Genotyping was performed using Sniffles2 with the --genotype-vcf option to assess whether SVs were homozygous or heterozygous in the original sequencing data.

SVs larger than 50 bp were extracted and classified for transposable element (TE) content. SV sequences were aligned to a comprehensive TE consensus library using minimap2 (--paf-no-hit). SVs were classified as TE-containing if ≥80% of the SV sequence aligned to TE consensus sequences. TE classifications were merged using bedtools to handle overlapping alignments, and SVs were categorized as containing single TEs, multiple TEs, or no TE content.

### Chromatin state analysis at TE insertions

Full-length TE insertions were identified by aligning TE consensus sequences to the genome assembly using BLAT [79]. High-confidence insertions were defined as those with near end-to-end alignment (query start <50 bp, query end >query size-50 bp) and ≥80% sequence identity. H3K9me3 ChIP-seq signal was analyzed around TE insertion sites using deepTools (v3.5.4) [95]. Signal matrices were computed using computeMatrix scale-regions with 5 kb flanking regions and 10 bp bins. Only TE insertions in euchromatic regions were analyzed, defined by chromosome coordinates excluding heterochromatic regions. Heatmaps were generated using plotHeatmap with k-means clustering (k=3) to identify distinct chromatin patterns. TEs were classified into “Piwi regulated” and “Piwi independent “ by a cutoff of >2.5x log2 fold-deregulation as determined in the RNAseq DGE analysis.

### RNAseq analysis

Illumina reads were processed by removing 3′ adapters with cutadapt [96], trimming the first five nucleotides, and filtering for minimal length (>18 nt) and sequence complexity (bbduk, entropy = 0.35, entropy window = 18, k = 4) [97]. Reads were first aligned to *Drosophila* rRNA precursor sequences and the mitochondrial genome using bowtie [83]. Remaining unmapped reads were mapped to the *OSC_r1.01* genome using STAR in two-pass mode with splice-junction support (--alignEndsType Local, --outFilterType BySJout, --alignSJoverhangMin 15, --alignSJDBoverhangMin 1, --outFilterMismatchNmax 1, --outFilterMultimapNmax 1000, --winAnchorMultimapNmax 2000) [80]. Alignments were separated into uniquely and all-mapping fractions. For visualization, HOMER [98] was used to generate tag directories (makeTagDirectory…- format bed-keepAll-single-fragLength given) and UCSC-compatible tracks (makeUCSCfile…-fsize 1e20-strand +/--fragLength given-noadj-normLength 0) were generated per strand. Counts were normalized to 10M uniquely aligned reads and final BigWig tracks were created with bedGraphToBigWig (Kent Utilities) [75], producing strand-specific (sense and antisense) coverage profiles for genome browser display.

For differential gene expression (DGE) analysis gene and transposon expression was first quantified by using Salmon [99] on a target file containing *Drosophila* reference transcripts and TE sequences in sense an antisense orientation (--dumpEqWeights -- seqBias --gcBias --useVBOpt --numBootstraps 100-l SF --incompatPrior 0.0 -- validateMappings). Salmon results were imported into R using tximport [100].

Gene-level counts were assembled into a DESeq2 [101] dataset with sample metadata, and genes with fewer than 10 total counts across all samples were filtered out.

Conditions were modeled using ∼ condition, and reference genotypes were defined by re-leveling as appropriate. Variance stabilization was applied using either variance stabilizing transformation (VST) or regularized log transformation (vst, blind = FALSE), depending on the number of detected genes. Differential expression analysis was conducted using DESeq2 with independent filtering, Benjamini–Hochberg correction (α = 0.05), and log2 fold change shrinkage with apeglm [102]. Additional gene-level GeTMM value statistics have been calculated using edgeR [103].

### ChIPseq analysis

Illumina reads were processed by removing 3′ adapters with cutadapt [96] and filtering for minimal length (>18 nt) and sequence complexity (bbduk, entropy = 0.35, entropy window = 18, k = 4) [97]. Reads were first aligned to *Drosophila* rRNA precursor sequences and the mitochondrial genome using bowtie. Remaining unmapped reads were mapped to the *OSC_r1.01* genome using bowtie [83] allowing for 1 mismatch.

Alignments were separated into uniquely and all-mapping fractions. For visualization, HOMER [98] was used to generate tag directories (makeTagDirectory…-format bed - keepAll-single-fragLength given) and UCSC-compatible tracks (makeUCSCfile-fsize 1e20-fragLength given-noadj-normLength 0). Counts were normalized to 10M sequenced reads and final BigWig tracks were created with bedGraphToBigWig (Kent Utilities) [75], coverage profiles for genome browser display.

### sRNAseq analysis

Illumina reads were processed by removing 3′ adapters with cutadapt [96], trimming 4 random nucleotides from each end and filtering for minimal length (>18 nt) and sequence complexity (bbduk, entropy = 0.35, entropy window = 18, k = 4) [97]. Reads were first aligned to *Drosophila* rRNA precursor sequences and the mitochondrial genome using bowtie [83]. Remaining unmapped reads were mapped to the OSC_r1.01 genome using bowtie allowing for 1 mismatch. Alignments were separated into uniquely and all-mapping fractions. For visualization, HOMER [98] was used to generate tag directories (makeTagDirectory…-format bed-keepAll-single-fragLength given) and UCSC-compatible tracks (makeUCSCfile…-fsize 1e20-strand +/--fragLength given - noadj-normLength 0) were generated per strand. Counts were normalized to 1M miRNAs and final BigWig tracks were created with bedGraphToBigWig (Kent Utilities) [75], producing strand-specific (sense and antisense) coverage profiles for genome browser display. For transposon histogram analysis, piRNA reads longer than 23 nucleotides were aligned to transposable element consensus sequences using bowtie (*--best--strata-f-v 2-a*), permitting multiple mappings. Alignments were subsequently filtered to retain reads that mapped either uniquely or at most twice to a single transposon consensus, to account for long terminal repeat (LTR) redundancy. Reads mapping twice were assigned a fractional count of 0.5 to each location.

### PROseq analysis

Paired end Illumina reads were processed by removing the first mate, clipping 5′ adapters with cutadapt from the 2^nd^ mate read [96], filtering for minimal length (>18 nt) and sequence complexity (bbduk, entropy = 0.35, entropy window = 18, k = 4) [97]. Reads were first aligned to *Drosophila* rRNA precursor sequences and the mitochondrial genome using bowtie. Remaining unmapped reads were mapped to the *OSC_r1.01* genome using STAR [80] in two-pass mode with splice-junction support (-- alignEndsType Local, --outFilterType BySJout, --alignSJoverhangMin 15, -- alignSJDBoverhangMin 1, --outFilterMismatchNmax 1, --outFilterMultimapNmax 1000, --winAnchorMultimapNmax 2000). Alignments were separated into uniquely and all-mapping fractions. For visualization, HOMER [98] was used to generate tag directories (makeTagDirectory…-format bed-keepAll-single-fragLength given) and UCSC-compatible tracks (makeUCSCfile…-fsize 1e20-strand +/--fragLength given - noadj-normLength 0) were generated per strand. Counts were normalized to 10M uniquely aligned reads and final BigWig tracks were created with bedGraphToBigWig (Kent Utilities) [75], producing strand-specific (sense and antisense) coverage profiles for genome browser display.

### *flamenco* splicing analysis

The OSC_r1.01 genome was indexed with STAR [80], and paired-end RNA-seq data were split into single reads, reoriented, mapped as single-end data with STAR using -- alignEndsType EndToEnd and two-pass mapping. Uniquely aligned reads were extracted with samtools (NH:i:1) [62] and indexed. Spliced alignments were identified from CIGAR strings containing “N” operators and converted to BED12 [63]. Splice junctions were collapsed with bedtools groupby, and junction counts were combined with flanking coverage to produce UCSC “interact” format tracks. Additional scoring was introduced by quantifying mean read coverage in a 1 nt upstream window of donor sites relative to junction read support. Junctions with at least three supporting reads and upstream coverage greater than ten were retained and scaled to UCSC scores (0– 1000). Final splice junction interact tracks were converted to BigBed with bedToBigBed using a custom interact.as schema, allowing direct genome browser visualization of putative splicing within *flamenco*.

### *flamenco* silencing by tethering - analysis

The initial 800 kb of the *flamenco* locus and additional control regions were subdivided into 100 bp non-overlapping tiles, and uniqueness was estimated from 25-mer mappability tracks (1 mismatch allowed). For each tile, uniquely mappable positions and percent unique coverage were calculated. In parallel, the entire X chromosome was partitioned into 1-kb tiles on both strands. Tiles were intersected with 25-mer uniqueness tracks, and only those with >50% uniquely mappable positions were retained. Small RNA alignments from individual libraries were then intersected with the tiled features, retaining only uniquely mapped reads. Counts were normalized per library to 1 million miRNAs. For each tile, normalized counts and counts per unique position were calculated, and final count tables were compiled across libraries to allow quantitative comparisons of occupancy along *flamenco*, additional control regions, and across the full X chromosome. Small RNA count tables derived from tiled genomic regions were imported into R. Because of their close similarity, the 6-and 7-day samples were treated as replicates. Tiles with ≤50% uniquely mappable positions and libraries outside the relevant time points were excluded. For fold-change analysis, tiles with insufficient coverage (≤1 count for locus-specific analysis or ≤75 counts for chromosome-wide analysis) were filtered out. Fold changes were then calculated by dividing normalized values of treatment samples (IDR-tethered) by their corresponding controls (GAL4).

### TE cluster coverage analysis

Transposable element consensus sequences were indexed using bowtie. Wild-type small RNA reads longer than 23 nucleotides were truncated to 25 nucleotides and aligned to TE consensus sequences using bowtie (-a-M 1--best –strata) [83].

Alignments were converted to BED format and strand-specific coverage was calculated using bedtools genomecov (-strand +/-) [62, 63]. For cluster coverage analysis, 25-mer sequences were extracted from piRNA cluster coordinates using bedtools getfasta and seqkit sliding (-s 1-W 25) for both OSC assembly and reference genome (dm6). These 25-mers were aligned to TE consensus sequences using bowtie (-v 1-a), allowing all valid alignments, and converted to strand-specific bedGraph format. Coverage tracks from small RNAs and cluster-derived sequences were merged using join to produce position-wise comparison tables. Coverage data were imported into R and converted to binary format using dynamic thresholds (cluster coverage >0, small RNA coverage >10% of mean TE coverage). Overlap scores were then calculated independently for sense and antisense strands. Specifically, for each TE and strand:

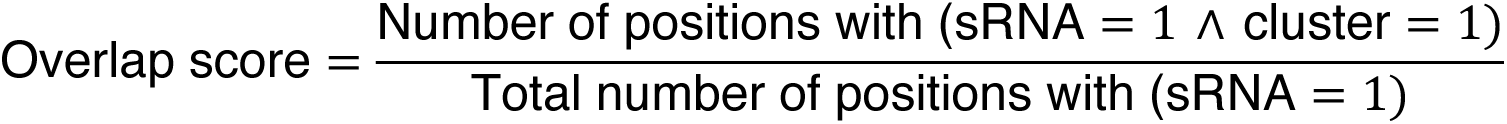

This value quantifies the fraction of small RNA–covered positions along a TE that are also represented by cluster-derived sequences. TE consensus elements with peak small RNA coverage <1000 reads or mean coverage <220 reads were excluded.

TEs with low peak coverage (<1000 reads) or mean coverage (<250 reads) were excluded from analysis.

### Statistical analysis and visualization

All statistical analyses and visualizations were performed in R (v4.5.1). Data wrangling and plotting were primarily carried out with the tidyverse collection (including *dplyr*, *ggplot2*) [104–106], with figure assembly using cowplot [107] and patchwork [108].

Interactive data exploration was supported by *plotly* [109] and formatted with the *scales* [110] package. Additional visualization packages included *ggbeeswarm* [111] *ggforce* [112], *ggh4x* [113], *khroma* [114], ggrastr [115] and *paletteer* [116] for custom themes, categorical palettes, and swarm/strip plots. JSON-based data handling employed *jsonlite* [117].

## DECLARATIONS

## Acknowledgements

We thank the Siomi labs for sharing the OSC line, the VBCF and IMBA/IMP/GMI core facilities for outstanding support, particularly the NGS facility for sequencing and the IT department for computational help. Members of the Brennecke laboratory gave valuable comments on the manuscript.

## Availability of data and materials

Genome assembly, Illumina and Nanopore genomic DNA sequencing, Hi-C, RNA-seq, small RNA, and PRO-seq data generated in this study have been deposited under BioProject PRJNA1338230 (pending GEO processing). Published datasets used in this manuscript include Nanopore direct RNAseq: PRJNA1236369 [118], STARR and DHS-seq: PRJNA175267 [119] & PRJNA248308 [120], Illumina RNA-seq: PRJNA724718 [53] and ChIP-seq: PRJNA178497 [121].

A detailed inventory of all datasets, including published data, is provided in Supplementary Table 1. The assembly is available via a UCSC Genome Browser hub with annotations and data tracks at https://brenneckelab.imba.oeaw.ac.at/Publication_Data/2025_Handler_OSC-genome/ and as a hosted UCSC session for direct browsing: https://genome-euro.ucsc.edu/s/Brennecke%2DLab/OSC_r1.01_Handler_et.al._2025. Code for genome assembly and analyses is available at https://github.com/BrenneckeLab/Handler_2025-OSC-genome.

## Author contributions

Conceptualization and writing: D.H. and J.B.

Data-generation and computational-analysis: D.H. Funding acquisition, resources and supervision: J.B.

## Consent for publication

Not applicable

## Ethics approval and consent to participate

Not applicable

