## Supplementary Material for "The Drosophila OSC Genome: A Resource for Studies of Transposon and piRNA Biology"

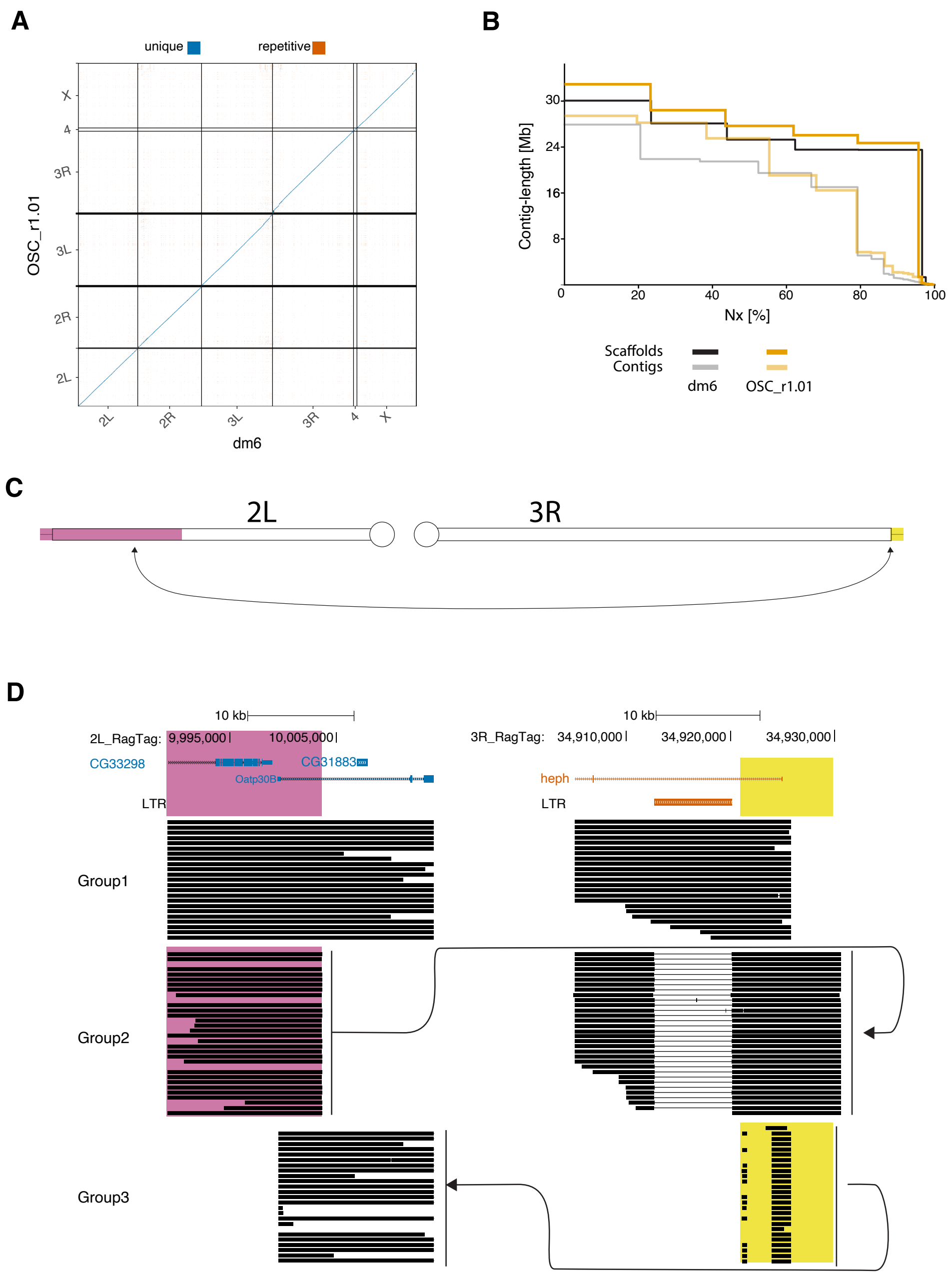

**Figure-supplement 1**

**(A)** Dot plots comparing the OSC genome with dm6, colored by alignment uniqueness. Alignments shorter than 5 kb were excluded. **(B)** Contig NG(x) plots of OSC and dm6 assemblies shown as scaffolds and as contigs after breaking at assembly gaps. **(C)** Schematic representation of the reciprocal translocation involving ~10 Mb from distal chromosome 2L and the distal end of chromosome 3R. **(D)** Nanopore read alignments confirming the heterozygous translocation between *CG33298* and *Oatp30B*. Group 1: reads supporting the non-translocated alleles. Group 2: reads spanning the translocated distal segment of chromosome 2L and the chromosome body of 3R. Group 3: reads spanning the chromosome body of 2L and the translocated distal 2L segment inserted into 3R. Reads in Groups 2 and 3 are aligned by sequence identity, with arrows indicating connections.

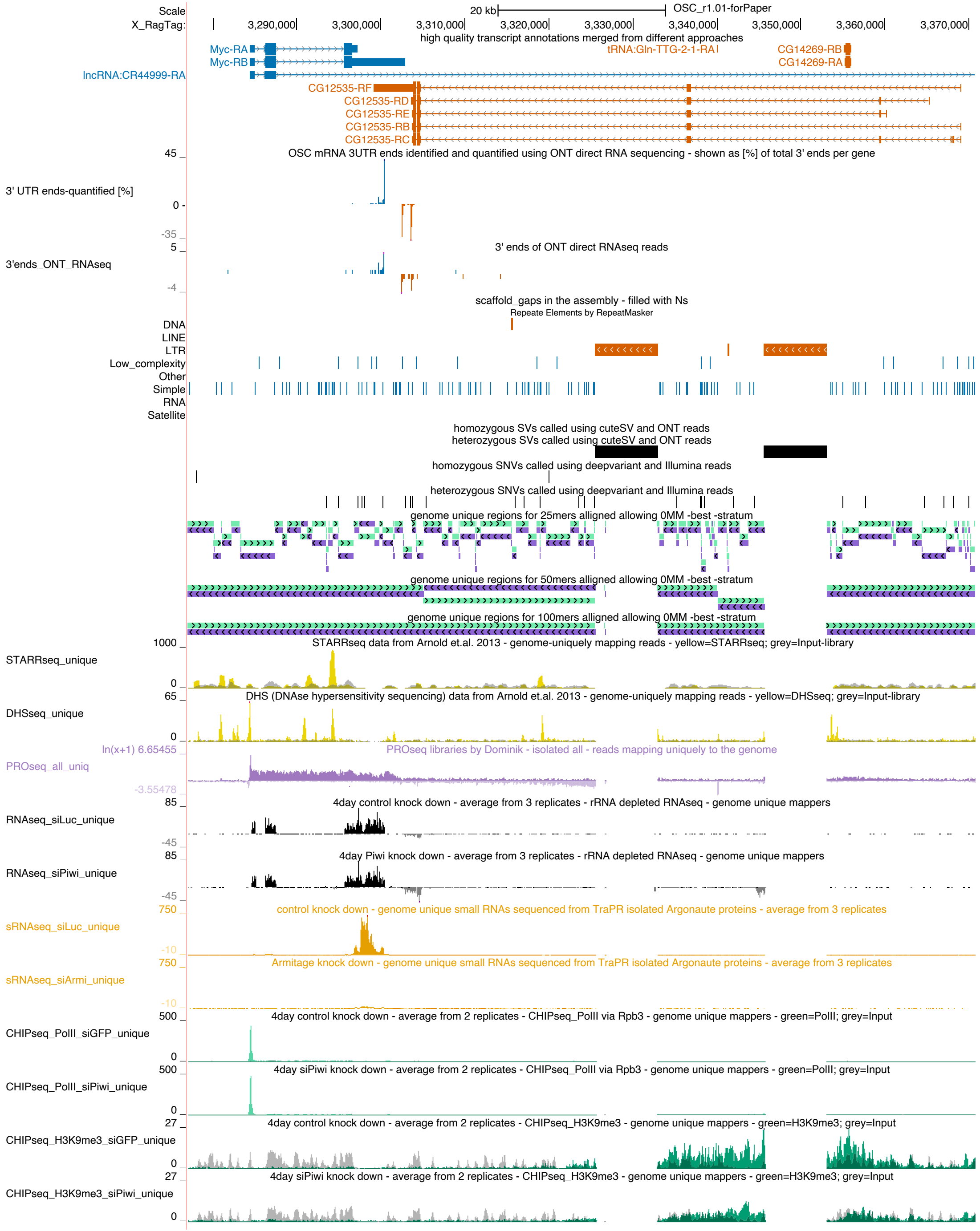

### Figure-supplement 2

Genome browser view of the *Myc* and *CG12535* locus as exported from the UCSC genome browser. (STARR, DHS, ChIP, PRO and RNA signal are shown as coverage per million reads, small RNA coverage normalized to 1M miRNA reads, data is displayed as the average of 3 replicates – PRO-seq signal is displayed as ln(x + 1)–transformed coverage). ChIP-seq is shown as an overlay of ChIP (green) and ChIP input (grey). STARR and DHS-seq are shown as an overlay of experiment (yellow) and input (grey).

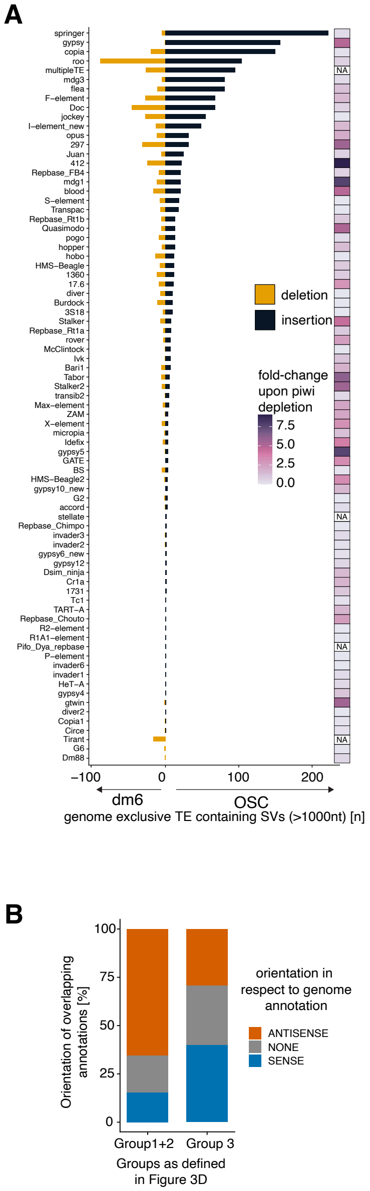

**Figure-supplement 3.**

(**A**) Structural variants (SVs) containing >80% transposable element content, shown per element. For each element, dm6-specific insertions are displayed on the left and OSC-specific insertions on the right. The accompanying heatmap indicates $\log_2$ fold change in expression upon *piwi* depletion, quantified from three replicate RNA-seq experiments (siControl vs. siPiwi). (**B**) Quantification of copia insertion orientation relative to overlapping expressed transcript orientation, stratified by H3K9me3 groups as in Figure 3D (with groups 1 and 2 merged into a single bar).

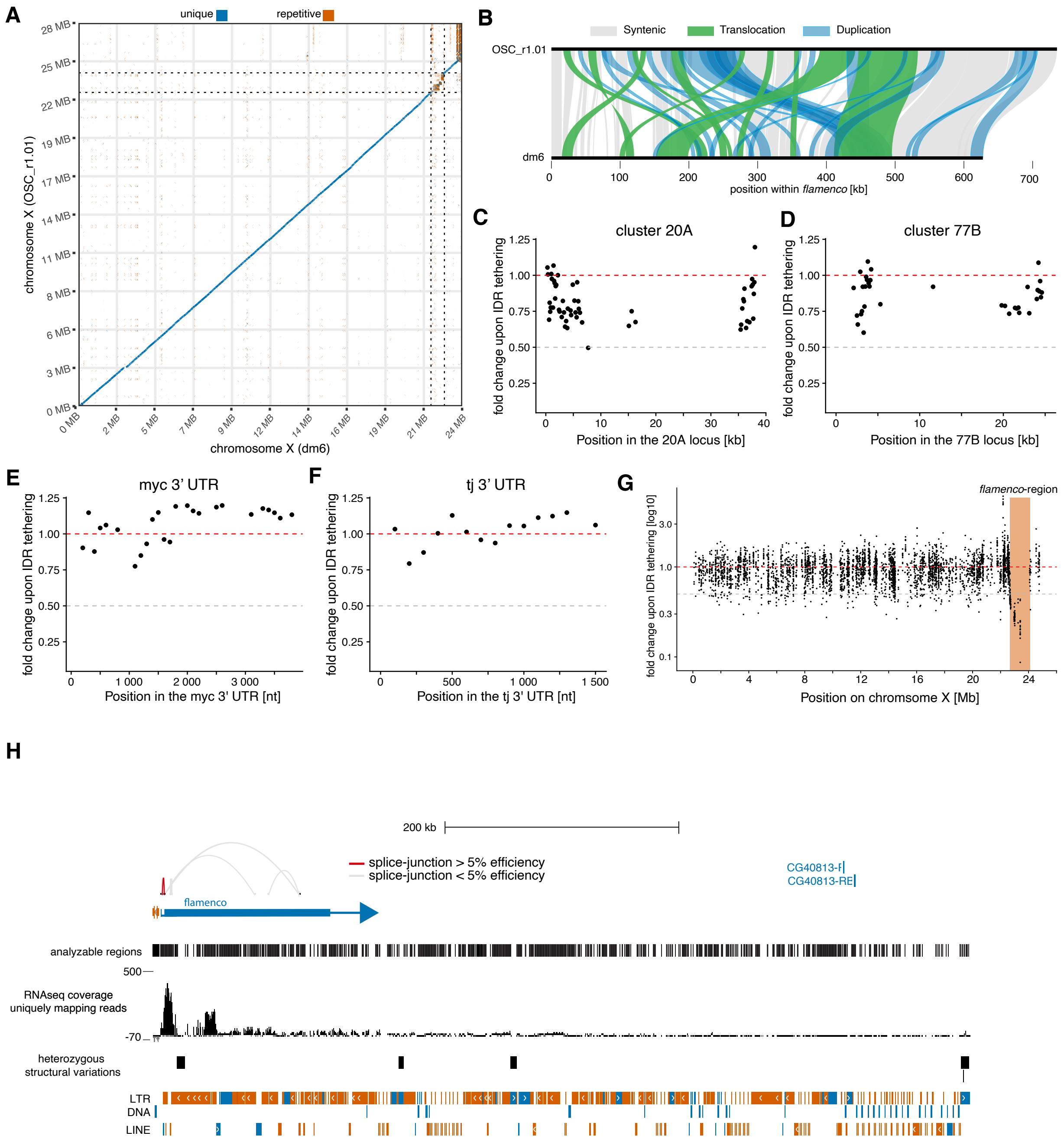

**Figure-supplement 4.**

(**A**) Dot plots comparing the X chromosome of the OSC genome with dm6, colored by alignment uniqueness. Alignments shorter than 5 kb were excluded. The extended *flamenco* locus spanning the region from *DIP1* to *CG14621* is indicated by dashed lines in both genomes. (**B**) Synteny map of the *flamenco* loci in the dm6 and OSC genomes. Syntenic regions, translocations, and duplications are highlighted in color; regions without synteny are left blank. (**C-F**) 1-kb tile analysis of fold change in small RNA levels upon repressor tethering to the *flamenco* promoter. Data are shown for *cluster 20A* (C), *cluster 77B* (D), the *myc* 3′ UTR (E), and the *tj* 3′ UTR (F). (**G**) 1-kb tile analysis of $log_{10}$ fold change in small RNA levels upon repressor tethering to the *flamenco* promoter, shown for the entire X chromosome. The extended *flamenco* locus spanning the region from *DIP1* to *CG14621* is indicated as a shaded area. (**H**) Genome browser view of the *flamenco* locus including the upstream *DIP1* gene. All identified splice junctions within the *flamenco* region are shown. Splicing efficiency was calculated as the percentage of spliced reads relative to the average read coverage within the preceding 5 nt. Analyzable regions are indicated, whereas regions excluded due to genome multi-mapping were not assessed for splice junctions. RNA-seq coverage of the reads used for splice analysis is shown as counts per million mapped reads.

**Supplementary Table 1**

| OSC genome assembly | Project-ID | GEO-ID | SRA-ID | Reference |
|---|---|---|---|---|
| Deposition of genome-assembly | PRJNA1338230 | | | this study |
| **Nanopore data** | | | | |
| siGFP_directRNAseq | PRJNA1236369 | GSE292040 | SRR32701027 | Voichek, M., et al. (2025) |
| siPiwi_directRNAseq | PRJNA1236369 | GSE292040 | SRR32701026 | Voichek, M., et al. (2025) |
| wtOSC_ONT_genDNA_RAD002 | PRJNA1338230 | | SRR35744053 | this study |
| wtOSC_ONT_genDNA_LSK108 | PRJNA1338230 | | SRR35744052 | this study |
| wtOSC_ONT_directRNAseq | PRJNA1338230 | | | this study |
| **DNAseq** | | | | |
| wtOSC_Illumina_genDNA | PRJNA1338230 | | SRR35744051 | this study |
| wtOSC_HiC | PRJNA1338230 | | | this study |
| STARRseq_Rep1 | PRJNA175267 | GSE40739 | SRR569903 | Arnold, C. D., et al. (2013) |
| STARRseq_Rep2 | PRJNA175267 | GSE40739 | SRR569904 | Arnold, C. D., et al. (2013) |
| STARRseq_Input | PRJNA175267 | GSE40739 | SRR569906 | Arnold, C. D., et al. (2013) |
| DHSseq_Rep1 | PRJNA175267 | GSE40739 | SRR569910 | Arnold, C. D., et al. (2013) |
| DHSseq_Rep2 | PRJNA175267 | GSE40739 | SRR569911 | Arnold, C. D., et al. (2013) |
| DHSseq_Input | PRJNA175267 | GSE40739 | SRR569912 | Arnold, C. D., et al. (2013) |
| hkCP_STARRseq_Rep1 | PRJNA248308 | GSE57876 | SRR1297291 | Zabidi M. A., et al. (2015) |
| hkCP_STARRseq_Rep2 | PRJNA248308 | GSE57876 | SRR1297292 | Zabidi M. A., et al. (2015) |
| hkCP_STARRseq_Input | PRJNA248308 | GSE57876 | SRR1297299 | Zabidi M. A., et al. (2015) |
| dCP_STARRseq_Rep1 | PRJNA248308 | GSE57876 | SRR1297296 | Zabidi M. A., et al. (2015) |
| dCP_STARRseq_Rep2 | PRJNA248308 | GSE57876 | SRR1297297 | Zabidi M. A., et al. (2015) |
| dCP_STARRseq_Input | PRJNA248308 | GSE57876 | SRR1297293 | Zabidi M. A., et al. (2015) |
| **RNAseq data** | | | | |
| siGFP_PolyA_Rep1 | PRJNA724718 | GSE173222 | SRR14311193 | Andreev, V. I., et al. (2022) |
| siGFP_PolyA_Rep2 | PRJNA724718 | GSE173222 | SRR14311194 | Andreev, V. I., et al. (2022) |
| siGFP_PolyA_Rep3 | PRJNA724718 | GSE173222 | SRR14311195 | Andreev, V. I., et al. (2022) |
| siPiwi_PolyA_Rep1 | PRJNA724718 | GSE173222 | SRR14311196 | Andreev, V. I., et al. (2022) |
| siPiwi_PolyA_Rep2 | PRJNA724718 | GSE173222 | SRR14311197 | Andreev, V. I., et al. (2022) |
| siPiwi_PolyA_Rep3 | PRJNA724718 | GSE173222 | SRR14311198 | Andreev, V. I., et al. (2022) |

| | | | | |
|---|---|---|---|---|
| wtOSC_rRNA-depleted | PRJNA1338230 | | | this study |
| **sRNAseq data** | | | | |
| siArmi_sRNA_Rep3 | PRJNA1338230 | | | this study |
| siArmi_sRNA_Rep2 | PRJNA1338230 | | | this study |
| siArmi_sRNA_Rep1 | PRJNA1338230 | | | this study |
| siLuc_sRNA_Rep3 | PRJNA1338230 | | | this study |
| siLuc_sRNA_Rep2 | PRJNA1338230 | | | this study |
| siLuc_sRNA_Rep1 | PRJNA1338230 | | | this study |
| flamenco-silencing_2xIDR-7days | PRJNA1338230 | | | this study |
| flamenco-silencing_2xIDR-6days | PRJNA1338230 | | | this study |
| flamenco-silencing_Gal4_1 | PRJNA1338230 | | | this study |
| flamenco-silencing_Gal4_2 | PRJNA1338230 | | | this study |
| **ChIPseq** | | | | |
| siGFP_H3K9me3_Rep1 | PRJNA178497 | GSE41729 | SRR609672 | Sienski, G., et al. (2012) |
| siGFP_H3K9me3_Rep2 | PRJNA178497 | GSE41729 | SRR609675 | Sienski, G., et al. (2012) |
| siGFP_Input_rep1 | PRJNA178497 | GSE41729 | SRR609678 | Sienski, G., et al. (2012) |
| siGFP_Input_rep2 | PRJNA178497 | GSE41729 | SRR609681 | Sienski, G., et al. (2012) |
| siGFP_Rpb3_Rep1 | PRJNA178497 | GSE41729 | SRR609684 | Sienski, G., et al. (2012) |
| siGFP_Rpb3_Rep2 | PRJNA178497 | GSE41729 | SRR609687 | Sienski, G., et al. (2012) |
| siPiwi_H3K9me3_Rep1 | PRJNA178497 | GSE41729 | SRR609674 | Sienski, G., et al. (2012) |
| siPiwi_H3K9me3_Rep2 | PRJNA178497 | GSE41729 | SRR609677 | Sienski, G., et al. (2012) |
| siPiwi_Input_rep1 | PRJNA178497 | GSE41729 | SRR609680 | Sienski, G., et al. (2012) |
| siPiwi_Input_rep2 | PRJNA178497 | GSE41729 | SRR609683 | Sienski, G., et al. (2012) |
| siPiwi_Rpb3_Rep1 | PRJNA178497 | GSE41729 | SRR609686 | Sienski, G., et al. (2012) |
| siPiwi_Rpb3_Rep2 | PRJNA178497 | GSE41729 | SRR609689 | Sienski, G., et al. (2012) |
| **PROseq** | | | | |
| PROseq-wtOSC_nuclei_biotin-C | PRJNA1338230 | | | this study |
| PROseq-wtOSC_permeabilized_biotin-C | PRJNA1338230 | | | this study |
| PROseq-wtOSC_nuclei_biotin-U | PRJNA1338230 | | | this study |

| | | | | |
|---|---|---|---|---|
| PROseq-wtOSC_permeabilized_biotin-U | PRJNA1338230 | | | this study |
| PROseq-wtOSC_nuclei_biotin-CU | PRJNA1338230 | | | this study |
| PROseq-wtOSC_permeabilized_biotin-CU | PRJNA1338230 | | | this study |
